# Slow ramping emerges from spontaneous fluctuations in spiking neural networks

**DOI:** 10.1101/2023.05.27.542589

**Authors:** Jake Gavenas, Ueli Rutishauser, Aaron Schurger, Uri Maoz

## Abstract

**Highlights:** 1. We reveal a mechanism for slow-ramping signals before spontaneous voluntary movements.

2. Slow synapses stabilize spontaneous fluctuations in spiking neural network.

3. We validate model predictions in human frontal cortical single-neuron recordings.

4. The model recreates the readiness potential in an EEG proxy signal.

5. Neurons that ramp together had correlated activity before ramping onset.

The capacity to initiate actions endogenously is critical for goal-directed behavior. Spontaneous voluntary actions are typically preceded by slow-ramping activity in medial frontal cortex that begins around two seconds before movement, which may reflect spontaneous fluctuations that influence action timing. However, the mechanisms by which these slow ramping signals emerge from single-neuron and network dynamics remain poorly understood. Here, we developed a spiking neural-network model that produces spontaneous slow ramping activity in single neurons and population activity with onsets ∼2 seconds before threshold crossings. A key prediction of our model is that neurons that ramp together have correlated firing patterns before ramping onset. We confirmed this model-derived hypothesis in a dataset of human single neuron recordings from medial frontal cortex. Our results suggest that slow ramping signals reflect bounded spontaneous fluctuations that emerge from quasi-winner-take-all dynamics in clustered networks that are temporally stabilized by slow-acting synapses.

## Introduction

Humans and other animals can initiate actions spontaneously, without immediate external triggers. Such spontaneous voluntary actions are critical to purposeful goal-directed behavior, but the neural mechanisms underlying these kinds of actions are poorly understood. Studies investigating the neural precursors of spontaneous voluntary actions have found that movement onset is preceded by neural signals that slowly ramp up or down more than 2 seconds before movement onset. This phenomenon is most evident in human medial frontal cortex and has been captured by scalp electroencephalography (EEG), where it is termed the readiness potential or “RP”^1–3^, as well as by fMRI^4–6^, and in single-neuron firing rates^7^, where it is termed the readiness discharge. Other mammals also exhibit slow-ramping signals before spontaneous actions in analogous brain regions^8–10^, and slow ramping has also been reported in invertebrates^11^. This suggests that slow ramping reflects an evolutionarily conserved mechanism related to voluntary action initiation. While widely reported, the origin and significance of slow ramping signals remain debated.

Slow-ramping signals were originally interpreted as reflecting preparatory processes that begin at the onset of ramping^2,12^. However, more recently, it has been proposed that slow ramping signals, like the RP, may instead be the result of autocorrelated fluctuations in neural activity that trigger movement upon crossing a threshold^13–17^ or bias the precise time of movement onset^18^. In contrast to classic interpretations, these models (hereafter collectively referred to as “stochastic fluctuation models” or SFMs) ascribe no special meaning to the onset of the slow-ramping signals. Rather, SFMs posit that ramping signals appear only because of back-averaging autocorrelated signals aligned to movement onset. SFMs therefore offer drastically different neural and cognitive implications for action initiation than classic interpretations^19^.

A major challenge for SFMs is that the time constant of empirically observed autocorrelations in neural spike trains is much shorter than the observed rate of ramping by about an order of magnitude. For example, slow ramping in the firing rate of human medial frontal cortex neurons evolves on the time scale of seconds^7^, but studies have reported that neurons in medial frontal cortex have autocorrelation time constants on the order of hundreds of milliseconds^20–22^ (although some have much longer time constants^23^). SFMs predict that more strongly autocorrelated fluctuations will approach the threshold more slowly and thus take longer to ramp up, while less autocorrelated (i.e., less temporally stable) processes are more likely to “jump up” and cross the threshold more abruptly, leading to shorter ramps^13^. It is thus unclear whether neurons that exhibit slow ramping are among the few that have longer autocorrelation time constants^23^, or increase their autocorrelation temporarily during slow ramping through some unknown mechanism, or if slow ramping is a network phenomenon that does not rely on the autocorrelation of single units.

Here, we utilized a combination of spiking neural network simulations and analysis of human single-neuron experimental data^7^ to investigate (1) the mechanisms by which temporally autocorrelated fluctuations in neural activity emerge in spiking neural networks, and (2) whether such fluctuations can account for the kind of slow ramping activity before spontaneous voluntary actions seen in the empirical data. We show that spiking neural networks can be configured to produce spontaneous fluctuations and slow-ramping activity with properties similar to those recorded^7^ from human medial frontal cortex during a spontaneous voluntary movement task similar to the classic Libet task ^2^. Our model is a spiking neural-network with 400 units with clustered connectivity. Such networks can exhibit spontaneous fluctuations^24,25^, but prior studies found that neurons in these networks exhibit “abrupt switching” activity—i.e. abrupt alternations between slow and fast firing—which is unlike the gradual changes seen during slow ramping.

We hypothesized that we could stabilize activity by inserting slow synapses, which are critical for producing temporally stable activity^26,27^ and increased autocorrelation in simulated spiking neurons^28^. We show that the presence of slow synapses led to more gradual spontaneous fluctuations and facilitated the emergence of slow ramping. The properties of ramping resembled the ramping signals observed in experimental data both at the single-neuron and population level. Furthermore, our model predicted that pairs of neurons that both ramped up or down would have more correlated activity compared to pairs that ramped in different directions even before the onset of ramping. This prediction was validated in the experimental data. We conclude that slow ramping signals before spontaneous, voluntary action likely emerge from backward-averaged spontaneous fluctuations aligned to threshold-crossings, and that clustered connectivity and slow synapses interact to jointly facilitate temporally smooth fluctuations. These results are compatible with SFMs of spontaneous voluntary action generation.

## Results

### Slow synaptic transmission temporally stabilizes spontaneous fluctuations in simulated networks

We simulated networks consisting of N=400 spiking leaky integrate-and-fire neurons (see LIF Network Simulations in Methods); 80% of the neurons were excitatory, i.e., they provided only positive inputs to other neurons, while the remainder were inhibitory. Excitatory neurons were grouped into 4 clusters of 80 neurons each, such that within-cluster connections were more likely than between-cluster connections. Simulations of a range of network sizes (N=200, 300, 400, 500 neurons) and number of clusters (N_clusters_ = 3, 4, 5, 8) led to the same qualitative results, described below, suggesting that our main results do not strictly depend on our chosen network size and number of clusters. The extent of clustering was quantified by the parameter R_EE_ (clustering degree), which is equal to the ratio of the probability of a within-cluster connection to the probability of a between-cluster connection (within-cluster connections were also stronger compared to between-cluster connections, as in^24^; see Connectivity Matrix Construction in Methods). Inhibitory neurons were not clustered, and their connectivity probability with excitatory neurons did not depend on R_EE_^29^.

R_EE_ describes a continuum between homogenous networks without clustering (R_EE_ = 1) and winner-take-all networks with strong clustering (R_EE_ >> 1) (Fig. 1A). Homogenous networks with balanced excitation and inhibition exhibit asynchronous spiking activity without slow spontaneous fluctuations in firing rates^30^. Conversely, in winner-take-all networks, neurons in a single cluster dominate and suppress neurons in other clusters via lateral inhibition implemented by inhibitory neurons^31^. In between these two extremes, clustered networks (roughly 2 < R_EE_ < 5, depending on network size, number of clusters, and other factors) exhibit slow spontaneous fluctuations due to a combination of lateral inhibition and lateral excitation between clusters. Note that in our networks, spontaneous fluctuations were entirely due to internal interactions because external input to the network was fixed (besides noise).

**Figure 1.**
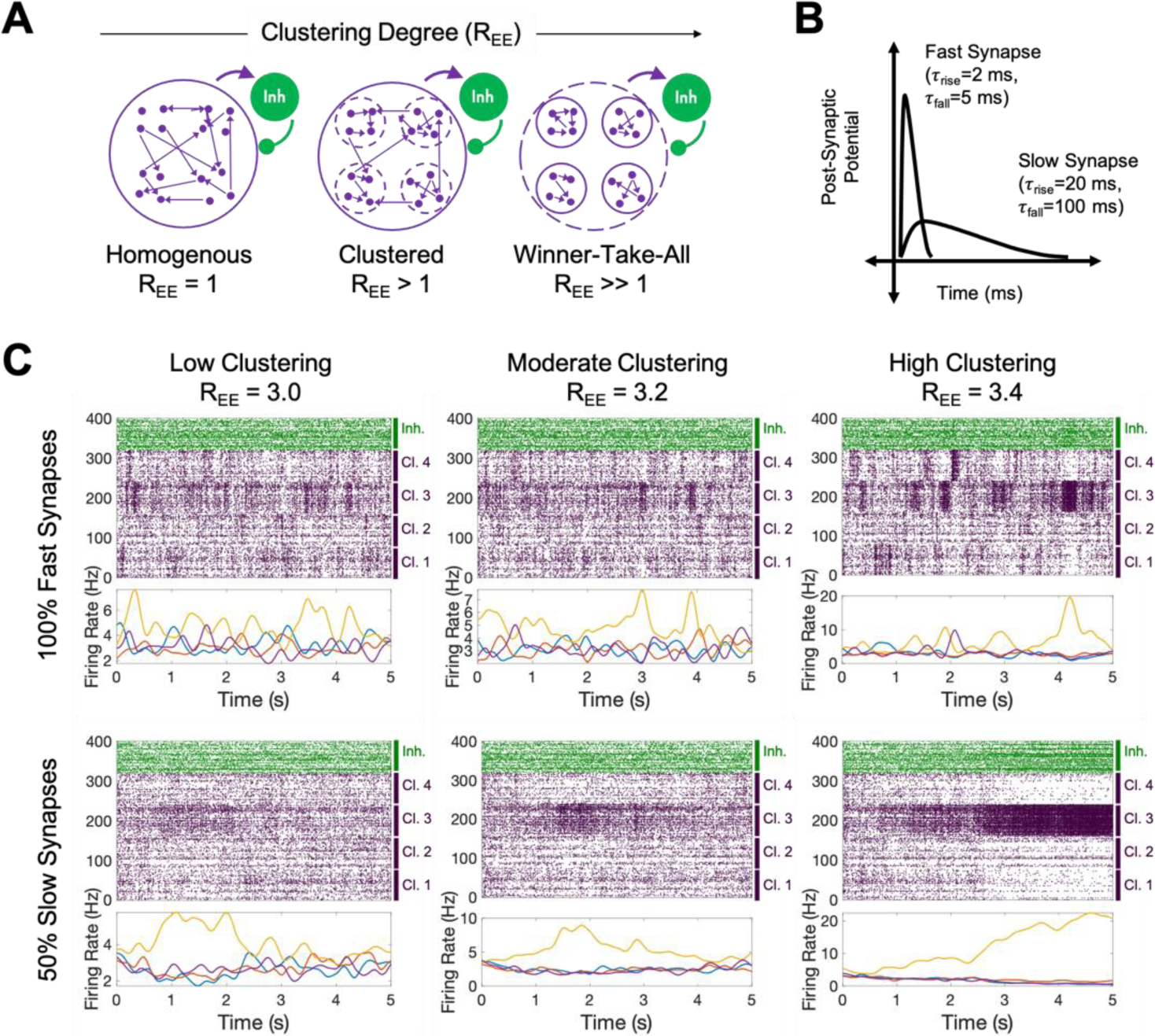
Simulation overview and example network activity. **A:** As the clustering degree (R_EE_) increases, connectivity transitions from homogenous/balanced, through clustered connectivity, to winner-take-all. **B:** Schematic of fast vs. slow synapses. **C:** Example 5-second simulations for networks with 100% fast synapses (top row) and 50% slow synapses (bottom row) at various clustering degrees (same underlying network structure, leading to the same ‘winning’ cluster, depicted in orange). For each subpanel, at the top are single-trial raster plots and at the bottom are cluster-averaged firing rates over time (spikes smoothed with 400 ms Gaussian kernel then averaged across all neurons in each cluster in different colors). In raster plots, purple are excitatory neurons (4 clusters of 80 neurons), green are inhibitory neurons. Clustering leads to spontaneous fluctuations, but fluctuation characteristics depend on synaptic dynamics. Networks with 100% fast synapses (top) switch between high-and low-activity states, whereas networks with 50% slow synapses (bottom) show smoother, more temporally stable fluctuations.

Previous work had only simulated clustered networks with fast synaptic dynamics, in which the entire postsynaptic potential is delivered within ∼5 ms^24,25,32^. We were interested in the role of slow synapses in modulating spontaneous fluctuations in such networks. We therefore simulated networks with both fast and slow synapses. We used double-exponential functions to model synapses^24,33,34^, with fast synapses having rising and falling time-constants of 2 and 5 ms, respectively, and slow synapses having rising and falling time-constants of 20 and 100 ms respectively^33^ (Fig. 1B). Because different types of synapses engage either fast or slow dynamics (AMPA and GABA-A for fast, NMDA and GABA-B for slow^34^), we modeled mixtures of synapses as a weighted sum of these two types of synapses, parameterized as the ratio of fast to slow synaptic signaling (i.e., the percent of post-synaptic potential (PSP) delivered via fast versus slow synapses; note that this applied to both excitatory and inhibitory connections; see LIF Network Simulations in Methods).

Recreating prior results for clustered networks with 100% fast synapses, we found that increasing R_EE_ was associated with a transition from asynchronous activity to more winner-take-all dynamics, with networks in an intermediate range exhibiting slow fluctuations (Fig. 1C top row). Notably, fluctuations in each cluster manifested as bistable switching activity, with entire clusters switching between states of low and high activity^24,25^. These fluctuations exhibited quasi-winner-take-all dynamics: activations in one cluster were accompanied by decreased firing in all the other clusters. Notably, switches between these states were rapid with no apparent slow ramping (Fig. 1C, top row; see Fig. 2 for quantification).

**Figure 2.**
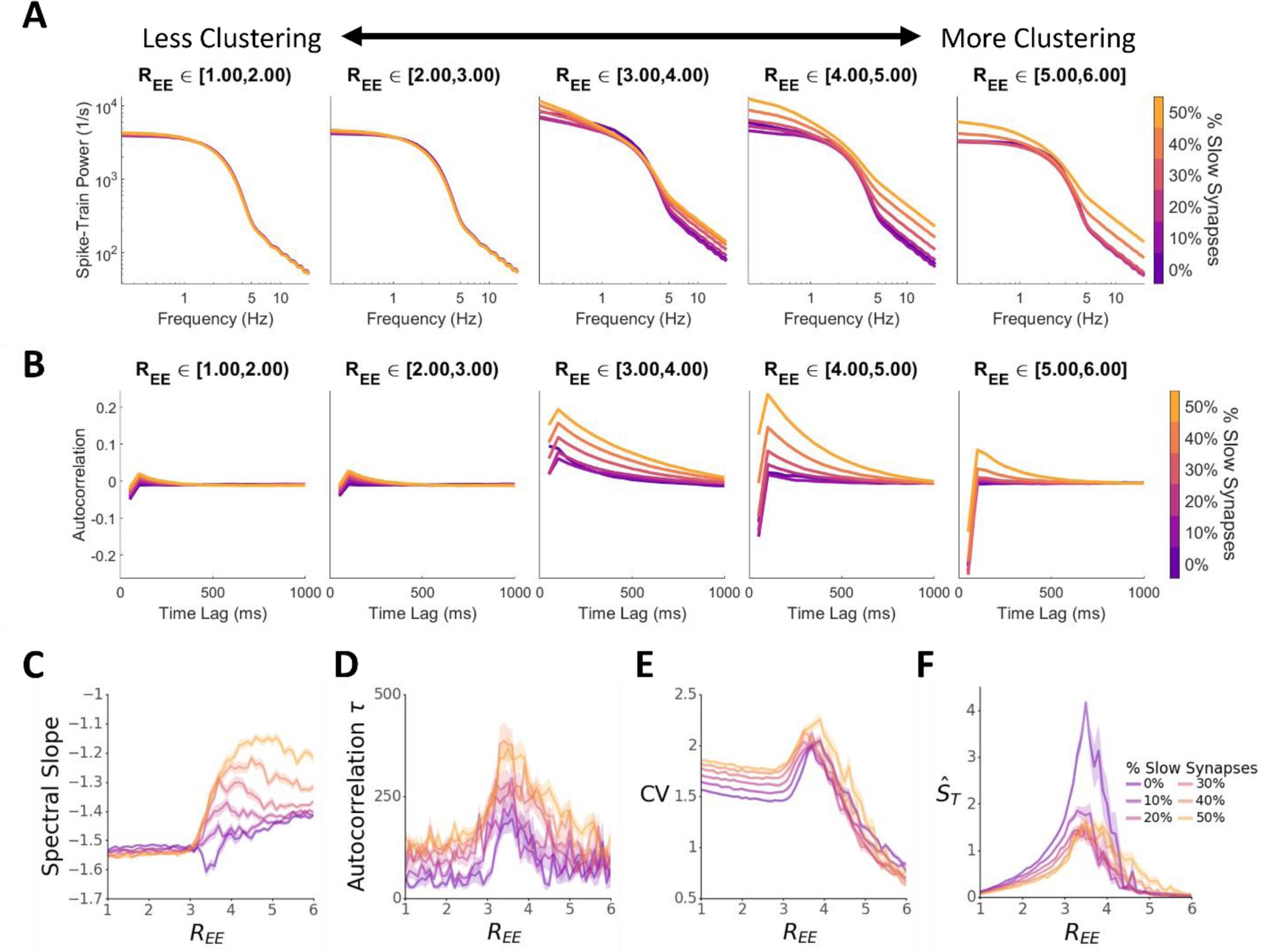
Effects of clustering & synaptic dynamics on temporal dynamics of neuronal activity. **A:** Effect of clustering and synaptic dynamics power spectra of simulated neurons’ firing rates (plotted in log-log scale). **B:** Effect of clustering and synaptic dynamics on spike train autocorrelation at time lags up to 1000 ms. **C-F:** Effect of clustering and synaptic dynamics on four measures of temporal dynamics. **C:** Spectral Slope, the slope of a linear regression of power to frequency in log-log scale (panel A **D:** Autocorrelation timescales, the decay parameter of exponential fit to autocorrelation functions (panel B). **E:** Coefficient of variation (CV), the firing rate standard deviation across time (100 ms non-overlapping bins) divided by trial-average firing rate. **F:** ŜT, a measure of firing rate variability across time like CV. ŜT was designed to specifically index slow-switching behavior like that seen in 100% fast / 0% slow synapse networks^25^.

In contrast, networks with slow synapses exhibited more gradual fluctuations. With 50% slow synapses, fluctuations in networks with intermediate R_EE_ values were temporally stable, with rises and falls that occur over a longer timespan when compared to networks with 100% fast synapses (Fig. 1C bottom row; see below for quantification). Fluctuations in 50% slow-synapse networks still exhibited quasi-winner-take-all dynamics, with activations in one cluster suppressing other clusters, but without the all-or-none nature of fluctuations common in 100% fast-synapse networks. Note that fluctuations are less clear in the inhibitory pool (Fig. 1C green raster); henceforth we exclusively analyze properties of the excitatory neurons.

### Network architecture and synaptic dynamics interact to shape network activity

Next, we quantified the effects of clustering degree and slow synaptic transmission on spontaneous fluctuations. We simulated 5 seconds of activity in 50 different randomly generated networks (each with 400 neurons, 320 excitatory) for R_EE_ values between 1 and 6 (increment of 0.1), with synaptic ratios ranging from 0% slow (100% fast) to 50% slow (increment of 10%, overall N=15,300 networks). We then investigated the effects of R_EE_ and synaptic ratio on measures of the networks’ temporal statistics, starting with power spectra. For each network, we calculated firing rate as a function of time for each neuron by smoothing spike trains with a gaussian filter (50 ms kernel), and then calculated the power spectra from the resulting smoothed firing rates (see Simulating & Analyzing Fluctuations in Methods). Next, we averaged power at matched frequencies across neurons within a network, and then averaged across networks within bins of increasing R_EE_ values (e.g. between 1.0 and 2.0; Fig. 2A left panel). Spectral power was generally higher at low frequencies and, on average, dropped monotonically as a function of frequency. As clustering increased past a critical value, spectral power increased for all networks. Notably, for low clustering coefficients (R_EE_ approximately between 1 and 3), the ratio of fast to slow synapses had no or a low impact on power. However, as clustering reached a critical range (R_EE_ above ∼3.0), a larger percentage of slow synapses resulted in higher power at relatively low (below ∼1 Hz) and high (above ∼5 Hz) frequencies. At even higher clustering (R_EE_ above ∼5.0), power decreased again, likely signifying the transition from the “fluctuation” regime to the “winner take all” regime.

We next investigated autocorrelation directly, employing the same grouping strategy as above (Fig. 2B; see Simulating & Analyzing Fluctuations in Methods). Unlike power, which did not vary as a function of synaptic dynamics for low clustering values, autocorrelation at short time lags was slightly higher for networks with a higher ratio of slow synapses. Then, at higher clustering values (R_EE_ above ∼3.0, roughly commensurate with results from power spectra), higher ratios of slow synapses led to noticeably greater autocorrelation at longer time-lags. As clustering increased further, autocorrelation at longer time-lags decreased and lag-one autocorrelation (corresponding to one 50 ms bin) became negative.

To further quantify the slow fluctuations we observed, we calculated four measures of spiking variability. First, we calculated the spectral slope (equal to the negative 1/f exponent) of neural firing rate power spectra as the slope of a linear fit of frequency to power in log-log space^13,35^ (Fig. 2C.; power spectra calculated as above; see Simulating & Analyzing Fluctuations in Methods). For low clustering (R_EE_ below ∼3.0), spectral slope did not depend on clustering degree and only little on the ratio of slow synapses, with higher ratios of slow synapses having slightly lower slope. At higher clustering degree, the dynamics changed, with spectral slope generally increasing with clustering, with larger increases for networks that had higher ratios of slow synapses. The higher slopes for networks with greater frequency of slow synapses indicates that power increases due to greater clustering were relatively larger at higher frequencies for higher ratios of slow synapses. Next, we calculated the autocorrelation timescale *τ*^20,21,36^ of neural firing rates as the decay parameter of an exponential fit to the autocorrelation function (Fig. 2D). Timescales were generally higher for networks with greater ratios of slow synapses for all clustering values. As clustering increased, autocorrelation timescales increased for all frequencies of slow synapses (until R_EE_∼3-4) and then dropped off. The drop-off was more gradual for networks with high percentages of slow synapses, indicating that slow synapses may impact the transition point from fluctuations to winner-take-all dynamics.

Next, we investigated how the coefficient of variation (CV) of individual neurons’ firing rates across time was affected by the clustering (Fig. 2E). The CV equals the standard deviation of firing rates across time (average firing rates in 100 ms non-overlapping bins; see Simulating & Analyzing Fluctuations in Methods), normalized by dividing by the trial-average firing rate.

High CV values are expected for networks that exhibit slow fluctuations in firing rates, as those fluctuations entail higher firing rate standard deviations across time. CV sharply increased after a critical R_EE_ value (∼3-3.5) for all networks, suggesting the emergence of slow fluctuations past that point. CV then decreased for higher R_EE_ values, indicating the transition to a winner-take-all regime. Notably, a greater percentage of slow synapses consistently resulted in a higher CV (though the rate of increase diminished with the proportion of slow synapses), suggesting that slow synapses increase firing rate variability even without clustering. We next looked at *Ŝ_T_*, a measure of spiking variability^25^ that characterizes slow-switching behavior in clustered networks, such as those seen in 100% fast synapse networks, Fig 1C top row). We investigated how *Ŝ_T_* was affected by clustering and slow synapses (Fig. 2F). *Ŝ_T_* is the standard deviation of cluster-averaged firing rates (100 ms non-overlapping bins) across time, averaged across clusters, and normalized by subtracting the same measure applied to random groupings of neurons rather than by cluster identity. Higher values of *Ŝ_T_* therefore increased temporal variability and slow-switching activity. *Ŝ_T_* increased steadily as R_EE_ increased, reaching maximal values for R_EE_ between 3 and 4, and then decreased. Notably, a greater percentage of slow synapses resulted in lower *Ŝ_T_* values. These results indicate that networks with slow synapses exhibit slow fluctuations, but that these fluctuations manifest as gradual fluctuations rather than slow-switching activity. Notably, however, the changes in temporal statistics we observed were heterogeneous across networks with the same R_EE_ and synaptic ratio parameters (especially so in the fluctuation regime; see Fig. S1), and likely depend on the emergence of specific connectivity patterns as clustering increases. Taken together, clustering is sufficient to lead to spontaneous slow fluctuations in spiking networks, yet our results suggest that introducing slow synaptic dynamics can greatly alter the nature of said fluctuations, in this case leading them to be temporally stable. We next turned to quantifying the speed of ramping in this network architecture.

### Slow-ramping at the single-neuron level

We next examined the relation between spontaneous fluctuations and slow-ramping signals at the single-neuron level. We simulated activity in 20 randomly connected networks with intermediate clustering degrees that led to spontaneous fluctuations (Min R_EE_ = 2.8; Max R_EE_ = 4; manually tuning for each network so that dynamics were in the fluctuation regime rather than winner-take-all regime; see Network Selection in Methods). For each network, we simulated 100 trials of 10 s each, when synaptic dynamics were either 100% fast or 50% slow. We then aligned data to times when activity in a specific cluster increased their activity above a threshold (termed threshold-crossing; the threshold was defined as 50% of maximum normalized firing rate across the whole trial; see Threshold Alignment in Methods). Neurons that significantly increased or decreased their firing rates in the 400 ms before threshold-crossing relative to a baseline (−3 to −2 s; as in Fried et al.^7^) were categorized as increasing or decreasing, respectively (Wilcoxon rank-sum test, the cutoff was p<0.01, as in Fried et al.^7^). The ratio of neurons categorized as increasing to decreasing was, on average, 50:50 in 100% fast synapse networks and 39:61 in 50% slow synapse networks.

Networks with 50% slow synapses demonstrated slow ramping with increasing and decreasing of firing rates before threshold-crossing, whereas networks with fast synapses showed only a steep jump shortly before threshold-crossing (Fig. 3A). Notably, our networks did not have a specified ‘ramping onset,’ nor were there any changes to the network’s input after trial onset (which was many seconds before the spontaneous ramping onset) beyond those attributable to noise.

**Figure 3:**
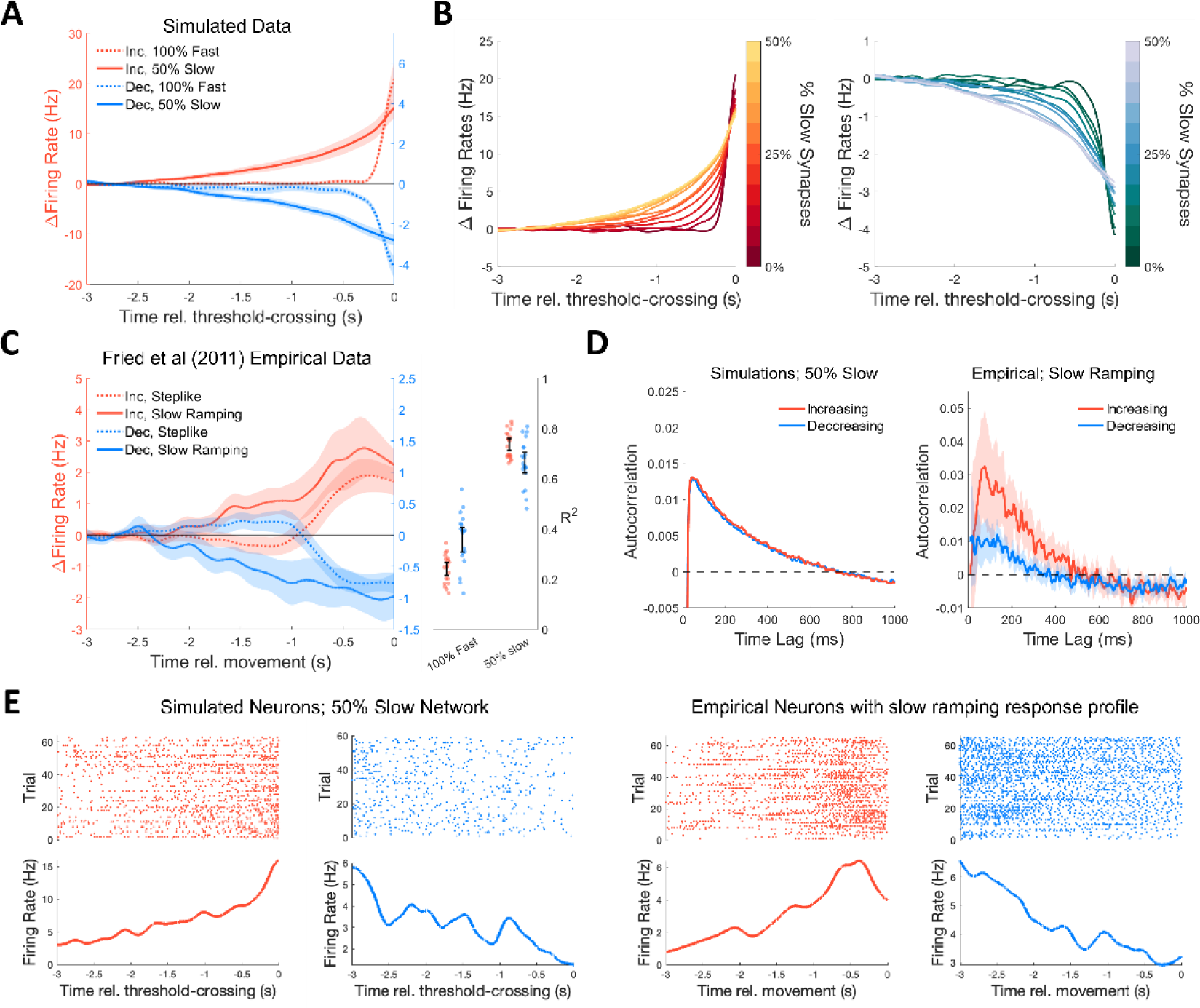
Slow ramping at the single-neuron level in simulated networks and recorded human frontal neurons. **A:** Firing rates (Δ, relative to baseline; mean ± 95% confidence intervals across networks) of neurons that increased (Inc.; red) or decreased (Dec.; blue) activity before threshold-crossings in networks with 100% fast (dotted lines) or 50% slow synapses (solid lines). **B:** Slow ramping of firing rates, either increasing (left) or decreasing (right), gradually emerges with the growing proportion of slow synapses. **C:** Left: Slow ramping before movement in human MFC neurons (Fried et al.^7^; Δ, relative to baseline; mean ± 95% confidence intervals across neurons). Some neurons show a steplike response profile (solid lines), and others show slow ramping (dashed lines). Right: Goodness-of-fit of simulated data versus experimental slow ramping response profiles (points: individual networks, error bars: 95% confidence intervals). For each network simulated with 0% and 50% slow synapses, we regressed average pre-threshold activity on the average empirical pre-movement activity at equivalent time-points (separately for increasing and decreasing neurons). From those models we extracted the R^2^ as a measure of how well each network fit pre-threshold changes in firing rates. 50% slow networks had significantly better fits compared to 100% fast networks for both increasing and decreasing neurons (p < 0.001 both increasing and decreasing). **D:** Slow ramping emerges despite low autocorrelation at the single-neuron level in both simulated (left; mean) and empirical neurons that exhibit slow ramping (right; mean ± 95% confidence interval). **E:** Example raster plots and trial-averaged firing rates from simulated (left) and empirical neurons (right). Notably, slow ramping is not apparent in most single-trial responses.

Therefore, the observed ramping must reflect the spontaneous fluctuations prior to threshold-crossing rather than a buildup driven by a specific intervening event. To further explore the relationship between slow synapses and ramping behavior, we conducted simulations (same as above) for various ratios of fast-to-slow synapses. Those simulations revealed that slow ramping emerged gradually as we increased the proportion of slow synaptic signaling (Fig. 3B). This gradual emergence closely matched the similar gradual emergence of slow ramping with increased autocorrelation (Fig. S2A). Accordingly, we found that increasing the ratio of slow synapses gradually increased activity autocorrelation during the ramping period at both the cluster and single-neuron level, although cluster-level autocorrelation was much higher than single-neuron autocorrelation (Fig. S2B). Together, these simulations support the notion that slow synapses facilitate temporally stable autocorrelated activity and, therefore, slow ramping.

### Comparison of single-neuron slow ramping in simulated and empirical data

We next assessed whether the degree of slow ramping observed in the model is comparable to that typically seen in the motor system preceding the onset of self-initiated actions. One of the key brain areas within which such slow ramping has been observed is in medial frontal cortex (MFC) of humans, including the pre-supplementary motor area, supplementary motor area proper, and anterior cingulate cortex^3,37,38^. We examined whether the properties of empirically measured ramping are comparable to those seen in our model by re-analyzing human MFC single-neuron data recorded by Fried and colleagues^7^ that show slow ramping and other firing rate changes before spontaneous voluntary actions.

We analyzed n=512 MFC neurons. Of these, 153 (30%) neurons significantly changed their response relative to movement onset (using the same selection criteria we used for our model data). Of these 153 neurons, 51 had increasing, and 102 had decreasing firing rates before movement onset relative to baseline. We next sub-selected neurons that exhibited slow ramping rather than more abrupt, step-like changes in firing rates. We distinguished between these two response profiles by fitting a sigmoid to trial-averaged firing rates and extracting an inverse-gain parameter. Neurons with an inverse-gain >0.4 were categorized as slow ramping, otherwise as steplike (see Neuron Categorization and Response Profiles in Methods). Thus, 41 MFC neurons were slow-ramping (27% of MFC neurons that significantly changed their firing rate before movement; 15 increasing and 26 decreasing) and the remaining 112 were steplike (73%; 36 increasing and 76 decreasing). Notably, slow ramping neurons showed progressive changes to their firing rates beginning around 2 s before movement onset on average, while steplike neurons showed a sharper firing rate change around 1 second before movement (Fig. 3C left; see Fig. S2C for distribution of ramping onset times). Here we focus on slow-ramping neurons.

Comparing the response profiles of simulated and recorded MFC neurons revealed that neurons in the slow ramping group had response profiles that were like those in simulated networks with slow synapses (compare Fig. 3A vs 3C). To quantify the goodness-of-fit of our networks, we regressed pre-threshold activity (averaged across neurons; −3 to 0 seconds rel. threshold-crossing) in simulated networks on the empirical data from matching time points (similarly averaged across neurons; Fig. 3C right). Networks with 50% slow synapses had higher R^2^ values than those with 100% fast synapses when comparing increasing neurons (mean [95% confidence interval]: 50% slow: 0.73 [0.71, 0.76] vs 100% fast: 0.24 [0.22, 0.27]; p < 0.001 two-sided sign-rank test) and decreasing neurons (50% slow: 0.66 [0.62, 0.71] vs 100% fast: 0.36 [0.31, 0.41]; p < 0.001 two-sided sign-rank test). Notably, slow ramping over ∼2 s at the single-neuron level occurred even though autocorrelation in these neurons decayed to zero after ∼600 ms, both for simulated and empirical neurons (Fig. 3D). Therefore, our model reproduces the key apparent paradox that we sought to resolve. Furthermore, visual inspection of raster plots from neurons that engaged in slow ramping behaviors showed that smooth, largely monotonic ramping was not visible on every individual trial—rather, activity was highly heterogeneous across trials (Fig. 3E).

### Slow ramping at the population-level

We next investigated slow ramping at the population level of all recorded/simulated neurons. We trained a linear discriminant analysis classifier (LDA) to differentiate activity at different points during the ramping period from a baseline (−3.0 to −2.6 s). Classification accuracy on statistically independent data, not used for training (5-fold cross-validation) for simulated networks with 50% slow synapses reached 75% at 641±7 ms before threshold-crossing (mean ± standard error here and below, averaged across networks), while classification accuracy for simulated networks with 100% fast synapses only increased above 75% on average 31±30 ms before threshold-crossing (Fig. 4A shows results from 100 simulations on a single network; further examples are in Fig. S4A). Classification accuracy in the simulated networks with 50% slow synapses resembled the gradual rise in accuracy obtained by an analogous analysis of the Fried et al. (2011) dataset (Fig. 4A right), in which we pooled MFC neurons across participants to create a pseudo-population for decoding^39^. Decoding accuracy reached 75% at ∼1.2 s before movement onset, and reached ∼90% at movement onset, presumably because many empirical neurons show a sudden ‘step like’ change in their firing rates at around 0.5s before movement onset (see Fig. 3C). Our model does not contain such neurons. These results suggest that our simulations best capture the early, gradually increasing accuracy in decoding activity from baseline up to ∼0.5 s before movement onset (Fig. 4A, left vs. right plot).

**Figure 4:**
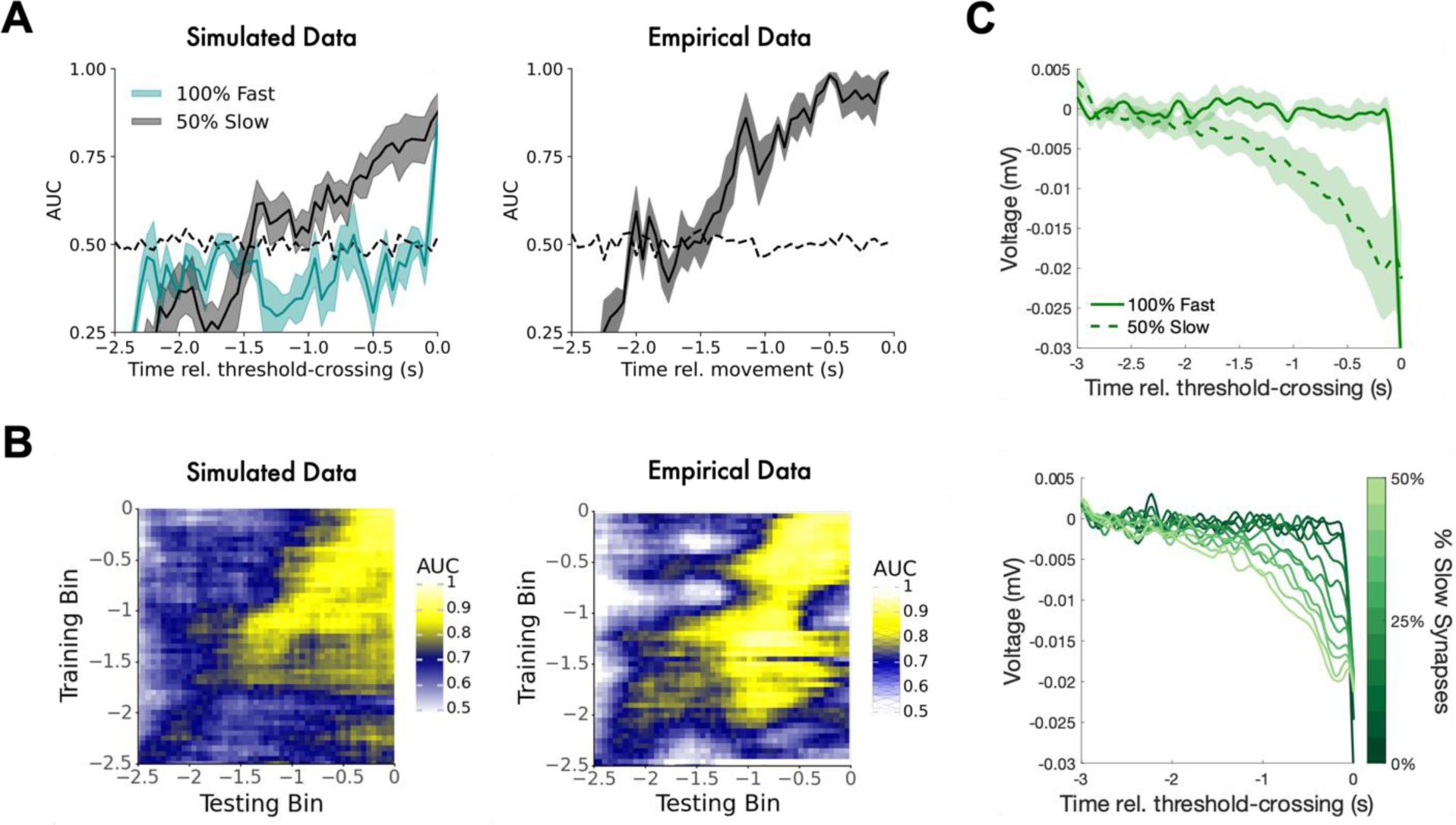
Slow ramping at the population level in simulated and empirical data. **A:** Classification accuracy versus baseline (linear discriminant analysis; spike count in 400 ms windows, times depicted refer to the leading edge of that window, versus −3 to −2.6 s baseline window) in simulated (left) and empirical data (right; shaded region is 95% confidence interval across cross-validations; dashed line is chance level across 100 shuffles of the data). Accuracy slowly ramps before threshold-crossing / movement, suggesting slow ramping at the population level. **B:** Performance of LDA classifiers trained at one time bin and tested on another time bin (relative to threshold-crossing or movement). Left: simulated data. Right: empirical data. Accuracy gradually increased leading up to the time of threshold-crossing or movement and was higher when the training bin was earlier than the testing bin compared to vice-versa (above versus below the main diagonal). **C:** Top: EEG proxy (PSPs onto excitatory populations) shows slow ramping in 50% slow but not 100% fast networks (confidence intervals over networks). Bottom: slow ramping in EEG proxy signal gradually emerges with an increased percentage of slow synaptic signaling.

To test whether gradually increasing accuracy reflected ramping at the population level, we investigated the temporal generalization of decoders, which can be used to ascertain the temporal evolution of neural activity^40^. We classified activity at different time points against a baseline (−3 to −2.6 s). If the population as a whole is exhibiting ramping activity, then earlier time points will be closer to the baseline in neuronal state space, while later time points will be farther from baseline. Therefore, when training a classifier on activity that is temporally close to baseline (early), the classifier will find a separatrix that is closer to the baseline in neural state space. Then, when applied to activity that is temporally further from baseline (later), the same separatrix will perform well because that activity is farther from baseline activity in state space (and vice-versa for training on later then generalizing to earlier activity). Thus, we should observe an asymmetry in across-time generalization accuracy such that classifiers perform better when generalizing from earlier to later timepoints compared to the reverse.

To test this theoretical prediction, we trained an LDA classifier to differentiate between ramping activity at a given time point relative to threshold crossing and activity during a baseline period (and tested that classifier’s accuracy at different time points (see Decoding Analyses in Methods). For simulated data, decoding accuracy gradually increased leading up to threshold-crossing (Fig. 4B left; more examples in Fig. S4B). Notably, as expected for ramping signals, across-time generalization accuracy was asymmetric—higher when training on earlier and testing on later timepoints compared to when training on later and testing on earlier timepoints (Fig. 4B). This can be seen by comparing the average values in the upper vs. lower triangle of the temporal generalization matrix (one-tailed rank-sum test p < 0.001). Similarly, for the MFC neurons, decoding accuracy showed a similar increase and asymmetry (rank-sum test comparing upper triangle to lower triangle decoding accuracies, p < 0.001, Fig. 4B right; see Fig. S4C for explicit demonstration of this asymmetry). To verify that this asymmetry reflects ramping, we generated spike trains that slowly ramped up and down to create a pseudo-population of neurons that we know a priori exhibit linear ramping (n=20, 50 trials, see Simulated Ramping in Methods). We conducted the same decoding and temporal generalization procedure on those spike trains and found the same asymmetry that decoders generalized to future time points better than to past time (one-tailed rank-sum test p < 0.001; Fig. S4D).

We also investigated ramping at the population level by calculating an EEG proxy from our network to compare to the readiness potential, a negative deflection in trial-averaged scalp EEG that slowly builds up over the last ∼1-2 s prior to movement onset^3,19^. Notably, we focus only on the early (negative) rise of the readiness potential, which is thought to originate in bilateral SMA, unlike the later parts, closer to movement onset, which originate in lateral motor cortex^3^. The EEG signal is typically taken to reflect the summation of post-synaptic potentials onto neurons^41^. So, as a proxy signal, we calculated the total signed post-synaptic input (both inhibitory and excitatory) delivered to each excitatory neuron in the network, then averaged across neurons, and investigated its trajectory leading up to threshold-crossings (see Threshold Alignment in Methods). Networks with 50% slow synapses exhibited slow negative ramping in the EEG proxy. In contrast, networks with 100% fast synapses had a relatively flat signal with an abrupt jump close to threshold-crossing (Fig. 4C top). Repeating this analysis for multiple ratios of fast-to-slow synaptic signaling revealed that slow ramping in the EEG proxy signal gradually emerged with a greater percentage of slow synapses (Fig 3C bottom). The results were similar for an EEG proxy that uses PSPs onto all simulated neurons rather than just excitatory neurons (Fig. S4E), accounting for suggestions that the human cortex contains larger proportions of inhibitory neurons than other species^42^.

### Pre-ramping noise correlations relate to slow-ramping activity

Our model, like other SFMs, predicts that slow ramping is a consequence of averaging autocorrelated fluctuations aligned to times of threshold-crossings. Those fluctuations do not begin at what would appear as ramping onset on average but occur throughout the entire trial (see Fig. 1C bottom). Therefore, a novel prediction of our model is that two neurons that show ramping behavior in the same direction (both increasing or decreasing) will have more correlated activity *before ramping onset* in comparison to pairs that show ramping behavior in different directions (one increasing and one decreasing). Such correlations in trial-by-trial firing rate are called noise correlations^43,44^, and are relevant to behavior^45,46^. Pre-ramping noise correlations that depend on the later type of ramping should emerge because ramping behavior is related to cluster membership—two neurons that both increase their firing rates are likely part of the same ‘winning’ cluster, and two neurons that both decrease their firing rates may be from any of the three ‘losing’ clusters, whereas two neurons that show opposite ramping directions almost certainly belong to different clusters (Fig. 5A).

**Figure 5:**
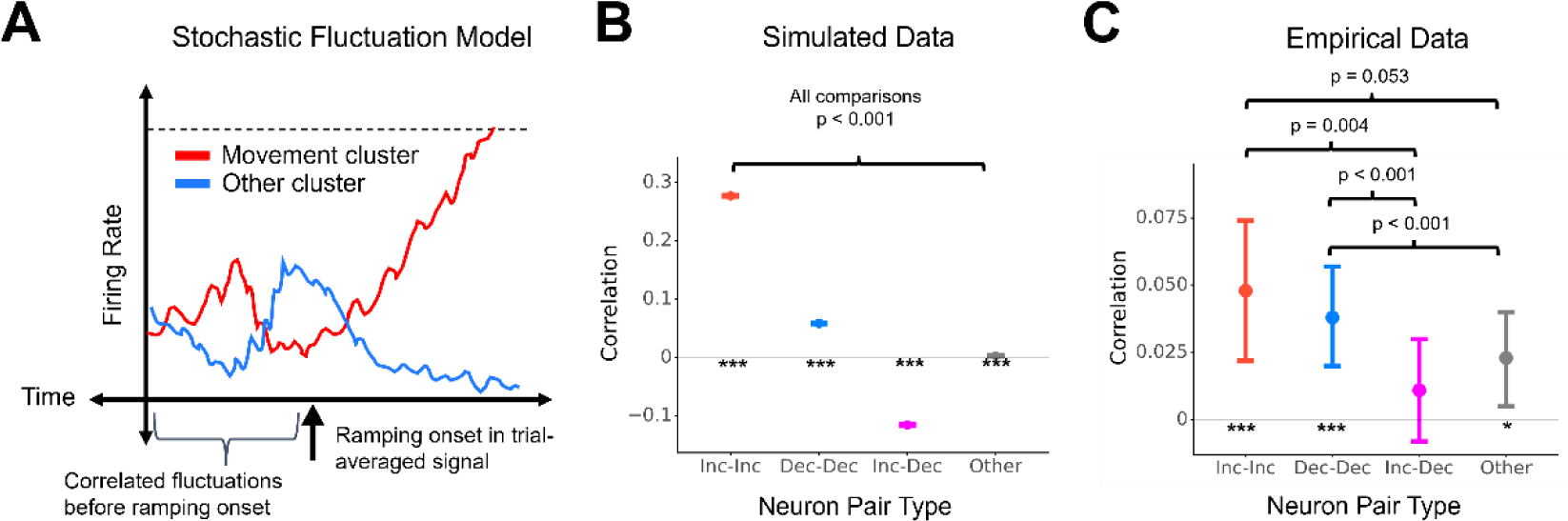
Model schematic and correlation structure relates to ramping direction. **A:** Stochastic fluctuation model (SFM) schematic. In SFMs, ramping reflects autocorrelated fluctuations prior to a threshold-crossing. Because the fluctuations occur throughout a trial, neurons with the same ramping behavior should exhibit correlated activity even before ramping onset. **B:** In 50% slow synapse networks, pairs of simulated neurons that exhibited the same ramping behavior—both increasing (Inc-Inc) or both decreasing (Dec-Dec)—were significantly correlated even before ramping onset (−3 to −2.5 s). Furthermore, neurons that showed opposite ramping behaviors (Inc-Dec) were anticorrelated before ramping. **C:** Pairs of empirical neurons that showed the same ramping behavior (increasing or decreasing) were significantly correlated before ramping onset (−3.5 to −2.5 s; mean & 95% confidence interval of pairwise correlation calculated via a linear mixed-effects model; steplike responses were categorized as “other” for this analysis). Pairs of neurons that exhibited the same type of ramping (both increasing or decreasing) were more correlated in the pre-ramping period compared to pairs that exhibited opposing types of ramping (increasing vs. decreasing). Although pairs showing opposite ramping behaviors were not reliably anticorrelated, as they were in the simulations, they were not statistically different from 0 in the positive or negative direction.

As predicted, in 50% slow synapse networks, activity in pairs of increasing neurons (i.e., both belonging to the specific cluster associated with movement) was positively correlated during a pre-ramping baseline period (−3 to −2.5 s, Fig. 5B, r = 0.277, p < 0.001, LME; see Analysis of Pairwise Correlations in Methods). In addition, activity in pairs of decreasing neurons (i.e., both belonging to clusters other than the one whose threshold-crossing determined movement timing) was also positively correlated, though to a lesser extent than pairs of increasing neurons, because they may belong to different clusters (r = 0.059, p < 0.001, LME). In contrast, activity in pairs involving one increasing and one decreasing neuron (i.e., very likely belonging to different clusters) was negatively correlated (r = −0.115, p < 0.001, LME). Finally, some neurons neither increased nor decreased their activity before threshold-crossing, likely due to the random nature of network connectivity leading to them receiving heterogeneous inputs. Activity in pairs of such neurons had very low but significantly positive correlations before ramping onset (r = 0.003, p < 0.001).

We next examined empirical noise correlations between firing rates in the pre-ramping baseline period (−3.5 to −2.5 s) for MFC neurons recorded in the same session (see Analysis of Pairwise Correlations in Methods; Fig. 5C). Pairs of neurons that later exhibited the same type of ramping (i.e., both increasing or both decreasing) were significantly positively correlated before ramping onset (increasing-increasing N=108 pairs: r = 0.048, 95% CI = [0.022 0.074], p < 0.001, LME; decreasing-decreasing N=865 pairs: r = 0.038, 95% CI = [0.020 0.057], p < 0.001, LME), whereas pairs with opposing types of ramping were not significantly correlated at baseline (N=532 pairs; r =0.011, 95% CI = [−0.008 0.030], p = 0.285, LME; Fig. 5C). Further, same-type pairs were more correlated than opposite-type pairs (increasing-increasing vs. increasing-decreasing: p = 0.004; decreasing-decreasing vs. increasing-decreasing: p < 0.001, LME post-hoc tests, Tukey correction for multiple comparisons) and were also more correlated than pairs involving neurons that did not increase or decrease or had a steplike response profile (N = 18,384 pairs; increasing-increasing vs. other: p = 0.053, only trending; decreasing-decreasing vs. other: p < 0.001). It is worth noting that noise correlations in the empirical data were smaller in magnitude overall compared to model predictions (although their magnitude is in line with prior studies^45,46^). Furthermore, noise correlations between opposite-type and other pairs were more positive compared to model predictions (e.g., opposite: r_simulated_ = −0.115, r_data_ = 0.011). Together, these results are in line with the predictions made by an SFM (Fig. 5A-B), increasing the plausibility that slow ramping signals reflect correlated fluctuations that are present from trial onset to threshold-crossing.

## Discussion

In this study, we demonstrated that spiking neural-networks can be configured to produce spontaneous fluctuations and slow-ramping activity with properties like those of intracortical experimental recordings in humans. Specifically, we recreated prior work that showed that networks with clustered architectures exhibit spontaneous “slow switching” fluctuations^24,25^, and further showed that slow synaptic dynamics temporally stabilized those fluctuations that (Figs. 1-2) and, in the aggregate, facilitated the emergence of gradual ramping activity before threshold-crossings (Fig. 3). In networks with 50% slow synapses, simulated neurons slowly ramped their firing rates, with 39% and 61% of slow ramping neurons increasing and decreasing their firing rates respectively. Interestingly, individual neurons in networks with 50% slow synapses slowly ramped their firing rates over 2 seconds before threshold-crossing, despite their autocorrelation dropping to zero at lags of ∼600 ms. By contrast, neurons in networks with 100% fast synapses did not exhibit changes in their trial-average firing rates until around 300 or 500 ms prior to movement in increasing and decreasing neurons respectively. Moreover, the accuracy of classifiers trained to distinguish activity from a baseline period slowly ramped up over time leading up to threshold-crossings (Fig. 4A left). Further, classifiers’ accuracy generalized to later time points better than they generalized to earlier time points (Fig. 4B left), indicating ramping at the population-level^40^.

Importantly, all of these features of simulated activity matched analogous analyses that we carried out on intracranially recorded human MFC neurons^7^ (Figs. 4A right, 4B right). Like simulated neurons, autocorrelation in real neurons that exhibited slow ramping diminished to zero within ∼600 milliseconds. In addition, an aggregate signal derived from our model, which could be interpreted as a proxy for EEG, slowly ramped negatively, akin to the readiness potential^1^ (RP) in terms of timing, shape, and ramping direction (Fig. 4C). Critically, all simulated slow-ramping signals emerged spontaneously from ongoing activity, without necessitating any external input at ramping onset to drive changes in activity. Finally, our model (similar to other SFMs) predicted that neurons that show the same ramping behavior (increase or decrease) after ramping onset would exhibit elevated pairwise correlations in their activity already before ramping onset. This prediction was borne out in the empirical MFC neurons, which exhibited a similar correlation structure to our model even before ramping onset (Fig. 5). This analysis was not a test of our model against classical interpretations of ramping onset (which suggest ramping onset reflects an event that drives ramping and, eventually, movement^2,12^, and thus make no particular prediction regarding pre-ramping correlations). But this qualitative confirmation of our model’s novel prediction in real data increases our model’s plausibility. Taken together, our study explains how highly autocorrelated spontaneous fluctuations can emerge in spiking neural networks through a combination of topographical and synaptic factors and demonstrates that such fluctuations are sufficient to explain several non-trivial aspects of slow ramping neural activity preceding spontaneous voluntary action.

One of our central aims was to investigate the discrepancy between short autocorrelation (decaying in ∼600 ms) and slow ramping (lasting ∼2 s—hence the period of slow ramping is therefore ∼3 times longer than the autocorrelation window for simulated and empirical neurons). However, slow ramping at the single-neuron level emerges only when averaging across trials, while autocorrelation is calculated from firing rates on individual trials. Inspection of raster plots from individual neurons that exhibit slow ramping (on average across trials) shows that smooth, generally monotonic ramping is not apparent on every trial (examples in Fig. 3E; more examples in Fig. S3; this fact is also noted by Schurger et al.^19^). Rather, activity on each trial is highly variable, and smoother, slow ramping emerges only in the trial-averaged firing rate. Our model gives a potential explanation: in our model, movement is triggered after an entire cluster’s average activity crosses a threshold (see Threshold Alignment in Methods). Therefore, individual neurons can and will show variable activity on individual trials. Still, trial-averaged response profiles will show smooth ramps because their activity is, across many trials, entrained to the activity of the cluster to which it belongs, which must increase (due to it later crossing the threshold) or decrease (due to lateral inhibition) its net firing before threshold-crossings.

Our work expands on research into spontaneous voluntary actions by offering what is, to our knowledge, the first model of slow ramping in spiking neural networks. Previous work had investigated spontaneous neural activity^24,25,28,30,32^ or autocorrelation in spiking neural networks^27,28,47^, but did not explore the connection to slow-ramping signals. Our model provides an implementation at the level of spiking neurons of previous, more abstract stochastic fluctuation models (SFMs) of spontaneous voluntary action, demonstrating how the fluctuations that are central to those models^13–17^ may emerge from biologically plausible spiking neural networks. Also, many brain regions are organized in a modular fashion^48^, and individual neural circuits have clustered connectivity among excitatory neurons^49,50^. Such clustering could plausibly emerge due to synaptic plasticity, such as Hebbian learning^51–53^, and ‘soft winner-take-all’ networks are hypothesized to be a canonical cortical circuit^54^. Furthermore, it is known that synaptic transmission occurs through both fast and slow channels^55^ (although, as is typical in computational modeling, our implementation of slow synaptic transmission is likely an oversimplification), and that some important cognitive functions rely on slow synapses^26^. Taken together, clustered connectivity and slow synaptic transmission are a plausible combination that we demonstrate can lead to spontaneous fluctuations and slow ramping. Additionally, the fact that our model can parsimoniously explain many features of MFC activity before spontaneous voluntary actions suggests that stochastic fluctuations in brain activity may underlie slow ramping signals and play a key role in the genesis of spontaneous voluntary action. Our model also goes beyond prior abstract computational models such as stochastic accumulation^14^ by describing a process that does not terminate at threshold crossing but is rather continuous in time. In doing so, our model offers possibilities for investigating strings or cascades of actions rather than single actions that occur in isolation.

Our results have major implications for the use of slow-ramping signals for predicting movement intentions, for instance in brain-computer interfaces. Because neural activity can stochastically increase or decrease, movement timing is not completely determined until threshold crossing. Therefore, slow-ramping signals may be of limited utility in online, real-time prediction systems. Indeed, predictions based on whole-brain EEG activity could predict movement timing above chance in real time only ∼600 ms before movements^56^ (see also^57,58^), whereas the RP and other slow ramping signals emerge 2 s or more before movement onset. Furthermore, in this respect, the EEG proxy we derived closely matched the RP. And our proxy is based on post-synaptic potentials, which exhibits a negative deflection before threshold-crossings due to an overall increase of inhibitory signaling and decrease of excitatory signaling as part of the quasi-winner-take-all process inherent to the networks we investigated. Taken together, we suggest that the apparent onset of the RP and other slow ramping signals several seconds before movement is the result of aligning spontaneous fluctuations to threshold-crossings rather than the many cognitive interpretations that have been offered since the RP’s discovery^3,19^.

### Future Directions

The model we developed is a “minimal plausible model” for slow ramping activity prior to spontaneous voluntary action. To our knowledge, it is the first model to describe how such ramping could arise spontaneously from spiking neural-network dynamics and captures several nontrivial aspects of premovement slow ramping activity. However, many questions remain about the neural mechanisms underlying spontaneous voluntary action. One key open question is regarding the timing of threshold-crossing and what mechanisms underlies its implementation.

Prior models suggested that threshold-crossing leads to movement within about 200 milliseconds^14^. However, many non-ramping neurons in MFC and other brain areas abruptly change their firing rates 0.5-1s before movement (Fig. 3C), around which time activity also abruptly changes in parietal cortex^59^. Such activations may reflect the spontaneous ignition of global activity patterns following threshold-crossings^17,60,61^. Future modeling work could investigate this by directly implementing a threshold mechanism and by recreating the activity of not just slow-ramping neurons but also those with a “step-like” response profile.

Slow ramping signals in the MFC have received much attention in the spontaneous voluntary action literature. Supporting this view, Fried and colleagues^7^ found that a greater fraction of neurons in MFC changed their activity prior to movement compared to temporal regions, suggesting a particular importance of this region. However, slow ramping signals have been observed in other regions, including medial temporal cortex^7^, sensory areas^6^, and subcortical areas such as the locus coeruleus (indexed by pupil dilations)^62^. Also, recordings in rats have found ramping activity in the dopaminergic substantia nigra^63–65^. Recent large-scale recordings of neural activity in mice further suggest that spontaneous voluntary movements involve widely distributed rather than localized activity^66,67^, as suggested by Schurger and Uithol^68^. Taken together, fluctuations influencing movement timing may reflect a distributed, brain-wide process rather than one localized only in MFC. One—admittedly speculative—possibility is that these fluctuations are related to fluctuations in arousal^69^, which may explain correlations between the RP and respiration^70^ and the presence of pupil dilations before freely timed movement in humans^62^. If so, our model might thus be implemented not in cortical regions exhibiting slow ramping activity, such as MFC, but rather in subcortical arousal hubs such as the Locus Coeruleus. Fluctuations and ramping in cortical regions could then be caused by input from these areas to the MFC^71^. Future studies should test this hypothesis by investigating the spatiotemporal relationship between slow ramping activity (including activity prior to the onset of slow ramping in trial-averaged activity, as we did in Fig. 5) and (1) peripheral signatures of arousal and activity in arousal related areas, such as the Locus Coeruleus, and (2) activity more proximal to the time of threshold-crossing or movement.

Crucially, because slow-ramping signals before spontaneous voluntary action may reflect stochastic population-level neural fluctuations, they also offer the opportunity to validate models that describe how temporally stable fluctuations emerge from neural dynamics. Brain activity fluctuates over several seconds or tens of seconds^72^, and the temporal properties of such fluctuations correlate with behavior^73^ change with age^74^, and are disrupted in disease^75^. It is therefore of interest to understand how these fluctuations emerge more generally. Indeed, prior work has also suggested that such fluctuations emerge when neural systems are poised at the edge of a phase transition^24,25,32,76–79^ (i.e., at criticality). In the present study, we manipulated the *clustering coefficient* of our networks to be just below a critical point, which had previously been shown to lead to spontaneous fluctuations in network activitys^24,25^. By introducing slow synaptic transmission, we showed that factors besides the parameter poised near criticality can alter the spatiotemporal dynamics of emergent fluctuations. Our model suggests that clustering and slow synapses lead to slow fluctuations emerging spontaneously in spiking networks (for instance, during resting periods). However, models using different mechanisms could potentially recreate our results. For instance, in our model we generated fluctuations by grouping neurons into clusters that had relatively higher (lower) intra-(inter-) cluster probability of connection between neurons. Schaub and colleagues^25^ found that other architectures (e.g. small-world connectivity) or varying intra-cluster synaptic strength instead of connection probability can also lead to spontaneous fluctuations. Furthermore, in our model, we temporally smoothed fluctuations via slow synaptic transmission. However, short-term plasticity or neuro-modulatory factors could potentially also stabilize fluctuations, and short-term plasticity could potentially alter the clustering degree, leading to further changes in fluctuation dynamics We elected to use slow synapses over these other possibilities due to their simplicity—slow synapses do not involve changes in connectivity, as short-term plasticity would, or factors external to the network, as neuromodulatory factors would, and thus our results emerge spontaneously. Future modeling efforts could explore whether combinations of these other factors would lead to results like ours and compare models to establish which factor, or combination of factors, best explain fluctuations in neural activity. Future studies could also investigate networks such as these analytically, as Schaub and colleagues did for clustered networks with fast synapses^25^, although we note that slow synapses might complicate such analysis by making network activity dependent on firing further in the past due to slower decay of synaptic currents (essentially making dynamics non-Markovian).

Slow ramping signals also emerge in other behavioral contexts, such as delayed response tasks (the contingent negative variation, or CNV, which is dissociated from the RP^80,81^), motor preparation^82,83^, and time perception^84,85^. Crucially, ramping in these other behavioral contexts occurs between two well-defined points—usually an external stimulus that leads to a response— whereas ramping onset in the spontaneous voluntary action context emerges without an external cue. So, whether slow ramping reflects similar processes across these contexts is unclear. In other “externally cued” contexts, reaction times are often too fast (hundreds of milliseconds) for “slow ramping” to occur beforehand, and even without time pressure to respond, pre-movement ramping, at least in MFC, is sometimes absent when decisions are not made spontaneously (e.g. according to stimulus value)^15^.

Interestingly, slow ramping with a seemingly spontaneous onset also occurs before other types of spontaneous behaviors, including generation of creative ideas^86^, spontaneous abstract decisions^87,88^, eureka moments in problem solving^89^, free recall of memories^79,90,91^, and switches between bistable percepts during binocular rivalry^92^. Notably, slow ramping signals before creative ideas have been linked to the temporal properties of spontaneous resting-state fluctuations^86^, similar to the relationship between autocorrelation and slow ramping^13^ that we investigated. Others have noticed these similarities and proposed that threshold-crossings by slow spontaneous fluctuations underlie various types of spontaneous behaviors. It is thus tempting to suggest that the neural mechanism we investigated in the present study may apply to spontaneous cognitive and perceptual processes more generally, offering an intriguing direction for future empirical studies.

## Acknowledgements

We thank Gabriel Kreiman for granting us access to the neural recordings we analyzed as part of this study and for discussion and feedback on an earlier version of this manuscript. We also thank Bill Newsome and Robert Kim for feedback on an earlier version of this manuscript. J.G., A.S., and U.M. were supported by the John Templeton Foundation and the Fetzer Institute (Consciousness and Free Will: A Joint Neuroscientific-Philosophical Investigation (John Templeton Foundation #61283; Fetzer Institute, Fetzer Memorial Trust #4189)). U.R. was supported by the US National Institutes of Health (U01NS117839) and J.G. and U.R. were supported by the NSF (BCS-2219800).

## Code and Data Availability

Code used for simulating data & analyzing simulated data will be made publicly available on OSF upon acceptance.

## Author Contributions

J.G., U.R., A.S., and U.M. conceptualized the project. J.G. conducted simulations and analysis of simulated and empirical data. U.R., A.S., and especially U.M. supervised and provided feedback on simulations and data analysis. J.G., U.R., A.S., and U.M. wrote the paper.

## Declaration of Interests

The authors declare no competing interests.

## Methods

### LIF Network Simulations

We simulated networks of leaky integrate-and-fire (LIF) neurons where the membrane potential *V*_*m*_ was governed (and updated at a resolution of 1 ms) by the equations:

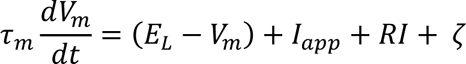

Here τ_*m*_is the synaptic time-constant (10 ms for our simulations). *E*_*L*_ is the reversal potential (−70 mV). *I*_*app*_ is a constant bias current set at the rheobase (300 pA). ζ ∼ *N*(0, 0.5 *mV*) is the noise term, which changes every 10 ms. In other words, each neuron receives constant noise for 10 ms, which then changes. The membrane potential, *V*_*m*_, had a threshold of −40 mV, and upon crossing it was reset to −65 mV and held constant for a refractory period of 2 ms. In every simulation *V*_*m* initial_∼ *U*(−70, −50 *mV*) for each neuron.

*RI* is the synaptic current vector. For each neuron it is a sum (weighted according to the connectivity matrix *W*; details below) of the other neurons’ synaptic output. Each such synaptic output is itself a weighted sum of fast and slow synaptic currents. *RI* follows the equation:

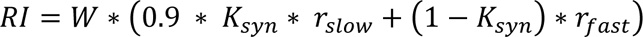

Here, 0 ≤ *K*_*syn*_ ≤ 1 is the ratio of fast to slow synapses. Hence, if *K*_*syn*_ = 0 or *K*_*syn*_ = 1 the network’s activity would entirely be governed by fast or slow synapses, respectively. And 0 < *K*_*syn*_ < 1, the network’s activity is governed by a weighted sum of fast and slow synaptic currents. We multiplied the slow synaptic currents by 0.9 to afford better comparisons between networks with similar clustering values (see below). Synaptic currents, *r*, follow a double-exponential time course including a rising and falling time-constant which differs between fast and slow synapses. Thus *r* follows the equations:

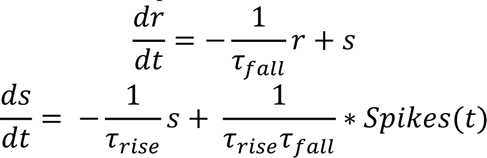

Here τ_*rise*_ and τ_*fall*_ are the time-constants for the synaptic currents to rise to their peak and then decay, respectively. Fast synapses had τ_*rise*_ = 2 *ms* and τ_*fall*_ = 5*ms* and slow synapses had τ_*rise*_ = 20 *ms* and τ_*fall*_ = 100 *ms* ^33^. *Spikes*(*t*) is a binary (0 or 1) spiking vector reflecting, for each time-step in the simulation, whether each neuron had spiked on the previous timestep. These spiking values were normalized by the rising and falling time constants, which ensured that the same total current was delivered for fast and slow synapses (see Fig. 1B).

### Connectivity Matrix Construction

Much of the dynamics of neural networks are determined by how neurons are interconnected, encoded in the connectivity matrix *W*. Previous research has found that *balanced* networks with roughly equal excitatory and inhibitory signaling exhibit stable asynchronous dynamics^30^. Balanced networks are usually implemented with more excitatory than inhibitory neurons (usually 75-80% excitatory), though with relatively weaker synaptic signaling in those excitatory neurons (to compensate for their higher proportion). To introduce slow changes in firing rates, we grouped the excitatory neurons into “clusters” both topographically and synaptically. Topographically, we raised the probability of connection for inter-cluster excitatory neurons in comparison with intra-cluster excitatory neurons. Synaptically, we increased the weights representing connections between excitatory neurons in the same cluster^24^ (although similar slow fluctuations in firing rates can be achieved by increasing intra-cluster connectivity strength alone, as well as through other methods^25^). Notably, however, the slow fluctuations in firing rates that emerged due to this clustering were depended significantly on the synaptic dynamics (see LIF Network Simulations section above).

We wanted to maintain sparse connectivity in our network, hence we used a 20% connection probability between any two excitatory neurons (*P*_*EE*_), taking into account the different number of intra- and inter-cluster connections. In a network with more than two clusters, there will be a larger percentage of inter-cluster connection opportunities than intra-cluster ones. So, calculating the precise values of the intra-cluster connection probability (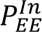) and inter-cluster connection probability (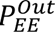) is non-trivial. To do so, we first note that the overall probability of connection (*P*_*EE*_) is a sum of intra- and inter-cluster connections divided by the total number of possible connections between excitatory neurons:

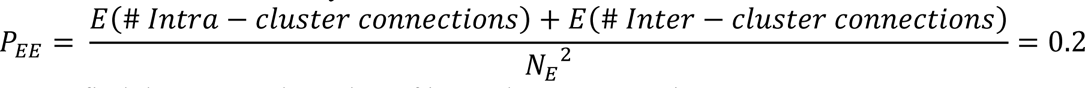

Next, we find the expected number of intra-cluster connections:

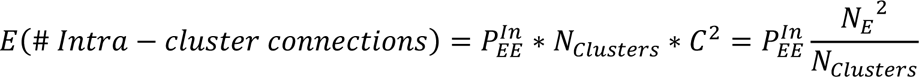

Where *N*_*clusters*_ is the number of clusters, *C* is the number of neurons in each cluster (which equals 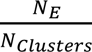), and *N*_E_ is the total number of excitatory neurons. The expected number of inter-cluster connections similarly follows:

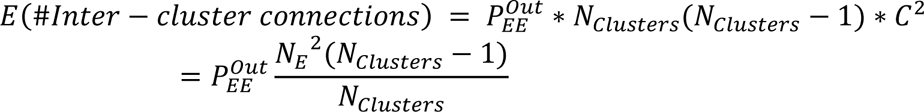

Then, letting 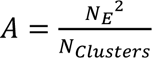 and 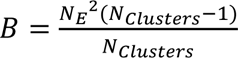 for simplicity, we obtain:

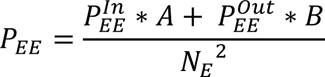

Finally, defining the clustering ratio R_EE_ as

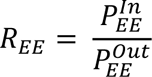

We can then solve explicitly for 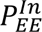 and 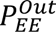:

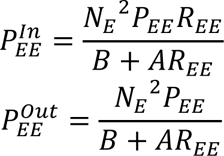

Solving for these probabilities explicitly thereby allows for a relatively easy construction of connectivity matrices corresponding to clustered networks, while maintaining a constant probability of connection overall (*P*_*EE*_=0.2, or 20% between excitatory neurons). Furthermore, these equations demonstrate that the precise values of 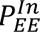 and 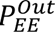 depend not only on the clustering coefficient (*R*_*EE*_), but also the number of excitatory neurons (*N*_*E*_), and the number of clusters (*N*_*Clusters*_) (or, equivalently, on the number of clusters and size of each cluster).

## Simulating & Analyzing Fluctuations

During simulations we varied two control parameters: the clustering coefficient, R_EE_ (defined below), and the ratio of slow-to-fast synapses, K_SYN_ (defined above). We simulated 5 s of activity in 50 unique network architectures for a range of R_EE_ values (1 to 6 at increments of 0.1) and a range of synapse ratios (0% to 50% slow synapses at increments of 10%). After discarding the first 500 ms of each simulation, we calculated several measures of temporal variability from simulated firing rates. First, we calculated the power spectra of firing rates (obtained by convolving spike trains with a gaussian kernel of width 400 ms) using the fast-Fourier transform in MATLAB and then taking the absolute value of the resulting complex-valued data. For calculating the spectral slope of these power spectra, we transformed data into log-log space and linearly regressed power on frequency, and then took the slope of the resulting linear fit as the spectral slope. For autocorrelation, we calculated firing rates using 50 ms non-overlapping bins and then calculated the autocorrelation of the resulting time-series up to lag 20 (corresponding to 1000 ms). For autocorrelation timescales, we fit an exponential function to the autocorrelation at lag *t* using the following equation:

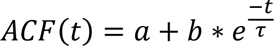

And took the decay constant (*τ*) as the autocorrelation timescale. We used the approach of Cavanagh and colleagues^22^ by fitting an exponential function to autocorrelation averaged across all excitatory neurons in a given network (as opposed to fitting and then averaging). We also followed prior research^20,22^ and excluded neurons with trial-average firing rates less than 2.5 Hz from further analysis, and omitted *τ* and autocorrelation of a network from further analyses if any of the following criteria were met: (1) fitted *τ* was not between 0 and 1000 ms, (2) fitted parameter *b* was below zero, or (3) if fitting of the exponential model did not converge. These rejection criteria led to omission of 3675 out of 15300 simulated trials (24%), but more than half of that (1913) came from networks with clustering coefficients above 4, at which point activity started transitioning into a winner-take-all regime.

Next, for CV, we calculated the firing rates of each excitatory neuron i ∈ {1,…, N}, where N is 320 (80% of 400), over time using non-overlapping bins of 100 ms. From these we calculated the coefficient of variation of firing rates across time (standard deviation divided by mean). For each neuron, with T time bins and firing rates at each bin we get a sequence Fi = [fi(t1), fi(t2),…, fi(tT)], over which we calculated the standard deviation σi and mean μi. Then, the network-level coefficient of variation is as follows:

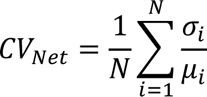

We calculated a single CV_Net_ value for each network, and plot grand means and standard error across networks in Fig. S1A. In Fig. S1B we plot individual network-average values (for the first 10 of the 50 networks to improve visibility).

In order to quantify spiking variability that was due to slow-switching or bistable activity, we also calculated the ŜT value used by Schaub and colleagues^25^ to investigate slow-switching behavior. This measure quantifies spiking variability over time by taking the average firing rate in each cluster i ∈ {1,…, C} in 100 ms non-overlapping bins. Then, for each cluster with firing rates in T time bins we get a sequence of firing rates F_i_ = [f_i_(t_1_), f_i_(t_2_),…, f_i_(t_T_)] where t_k_ represents the kth time bin, for which we calculate the standard deviation σ_T_(i), and then sum to obtain an initial measure of spiking variability.

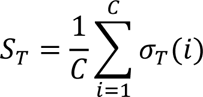

We then normalize this value by subtracting a bootstrapped value *S_Boot_* calculated on shuffled data (shuffling each neuron’s cluster membership), averaged over 20 random shuffles.

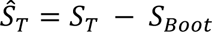

Like for CV, we calculated a single ŜT value for each network, and plot grand means and standard error across networks in Fig. S1C. For Fig. S1D we once again plot individual network-average values for the first ten of fifty networks.

### Network Selection for Threshold-Aligned Analyses

Prior work had shown that clustered networks exhibit fluctuations for a critical range of clustering coefficients (2 < R_EE_ < 5 typically, although this depends on network specifics; Litwin-Kumar & Doiron, 2012; Schaub et al., 2015). Because we generated network structures pseudo-randomly (setting the seed for random generation using MATLAB’s rng command), the critical range was slightly different for each network. We identified an R_EE_ value suitable for simulation in each network by visually inspecting activity at a range of values. We selected R_EE_ values for each network such that the network exhibited fluctuations at the given value for both 100% fast and 50% slow synapses, but exhibited winner-take-all behavior in at least one case if we increased the R_EE_ by 0.3. Prior work had linked the R_EE_ value demarking the transition to the winner-take-all regime to the leading eigenvalue of the connectivity matrix exceeding a value of 1 ^25^. However, our simulations did not exhibit such a relation, likely because that prior work used non-dimensionalized equations for simulating our activity whereas we used the dimensionalized versions of those equations.

### Threshold Alignment

We hypothesized that spontaneous fluctuations trigger movements upon crossing a threshold, leading to slow ramping in trial-averaged activity. To investigate this hypothesis, we simulated 10-second trials 100 times for each of the 20 network architectures and then aligned to threshold-crossings (defined below). We identified the most-active cluster across all simulated trials and aligned to threshold crossings in that cluster (reflecting the notion that threshold crossings in only some subpopulation of SMA would lead to a movement). If no threshold crossing was detected in a trial, the trial was omitted from further analyses.

Threshold crossings were defined as the time at which the most-active cluster crossed 50% of its maximum normalized firing rate across the entire simulation period. This was obtained by averaging firing rates across neurons in each cluster, subtracting the mean firing rate of the first 500 ms, and then dividing by the maximum firing rate on each trial. The choice to use a different threshold for each network was motivated by the fact that the magnitude and other features of fluctuations were heterogeneous across different networks even with similar clustering coefficients. Our choice of 50% of the maximum normalized firing rate was motivated by Fried and colleagues’^7^ finding that increasing neurons reached about 50% of their maximum normalized firing rate right before movement (Fig. 4 in Fried et al., 2011). Only trials where crossing occurred more than 3 s into the trial were retained for analysis of neural data (as is commonly done in self-initiated action paradigms^7^), as we were looking at changes across this period.

Neurons were categorized as increasing or decreasing by comparing their firing rates in the 400 ms before threshold crossings to their firing rates during a baseline period (−3 to −2 s, rank-sum test with a cut-off of p=0.01, as in^7^). Firing rates were smoothed with a gaussian kernel (width 400 ms), then averaged across response types (increasing and decreasing) to get a single trace for each network. Those traces were then averaged across networks for Fig. 2A; the shaded regions reflect the confidence intervals across networks.

Our EEG proxy was obtained by averaging the post-synaptic potentials (PSPs) onto excitatory neurons^41^. We retained the PSPs onto each neuron (*RI* in the above equations). We then sub-selected the PSPs to retain only those going onto excitatory neurons, averaging across neurons to obtain a single EEG proxy per trial. We then averaged across trials to get a single EEG proxy trace per network. Those traces were again averaged across networks for Fig. 2B, and the shaded regions reflect the confidence intervals across networks.

Autocorrelation at the single-neuron and cluster level was obtained using the autocorr function in MATLAB on the firing rates (smoothed using a moving average filter with a window of 10 ms to avoid smoothing artifacts; then averaged across neurons in each cluster for cluster-average firing rates) for each −3 to 0.5-second trial aligned to threshold-crossings. Those traces were then averaged across trials to get a single autocorrelation trace for each neuron or cluster. Then, those traces were averaged across neurons or clusters to get a single trace per network and then averaged across networks to obtain grand averages for Fig. 2C.

### Neuron Categorization & Response Profiles

We categorized empirical and simulated neurons as increasing or decreasing following the process described by Fried et al.^7^. We compared firing rates in the 400 ms before threshold crossing or movement to firing rates during a baseline period (−3 to −2 s) using a non-parametric rank-sum test. Neurons that significantly increased (decreased) their activity relative to baseline were categorized as increasing (decreasing; cut-off was p=0.01). Notably, this process omits neurons with more complex response profiles (e.g. Fig. 3 in Fried et al., 2011^7^), but these are out of the scope of this study.

MFC neurons had a mixture of response profiles. In particular, some neurons slowly ramped their firing rates up or down before movement, and others exhibited a steep jump up or down before movement. Simulated neurons tended to exhibit the same type of response (slow ramp or steplike), but responses in empirical data were mixed and therefore required a method to distinguish them. To do so, we followed a process similar to the supplementary analysis conducted by Fried et al.^7^ (supplementary analysis/Fig. S2 in their article). We fit a sigmoid function to the trial-averaged smoothed firing rates (400 ms Gaussian) using nonlinear least-squares via the Levenberg-Marquardt algorithm in MATLAB. We fit a different sigmoid depending on if the neuron was increasing or decreasing:

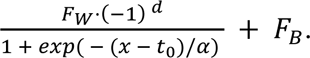

Here F_B_ is the firing rate at baseline, F_W_ is the firing rate in the 400 ms before movement. t_0_ is the (fitted) sigmoid inflection point. α is the (fitted) inverse-gain parameter (i.e. higher values of α correspond to a smoother ramp). And d is 0 for increasing neurons and 1 for decreasing ones. We selected an inverse-gain of 0.4 as the cut-off for steplike versus slow ramp response profiles based on visualizing averaged firing rates from the Fried et al.^7^ (2011) dataset (see Fig. 2C). Thus, neurons with an α parameter greater than 0.4 were categorized as slow-ramping, and neurons with an α parameter less than 0.4 were categorized as steplike.

### Decoding Analyses

We also assessed whether above-chance decoding accuracy in the seconds before movement^7,37^ might emerge from aligning fluctuations like those in our networks to threshold-crossings. For simulated data, at each time point, we compared spike counts in the time bin [t-0.4, t] s to spike counts in the baseline of (−3,−2.6) s for all neurons. We first constructed a pseudopopulation of neurons for which we had more than 40 trials by grouping them into one dataset for purposes of the decoding analysis. We trained a linear discriminant analysis classifier on 80% of the data and recorded its performance on a reserved 20% test set (i.e., 5-fold cross-validation). Fig. 3A shows the average accuracy and standard error across 5 cross-validation splits for individual networks (the same splits were used for *K*_*syn*_ = 0 and *K*_*syn*_ = 0.5), as well as performance on 100 random splits of the data to assess chance-level performance. For empirical data, we constructed a pseudo population of all recorded MFC neurons. We aggregated across subjects with at least 50 trials and conducted the same analysis as above, with time relative to movement onset rather than threshold-crossing.

To assess the temporal generalization of decoding (Fig. 3B), we completed a similar analysis as above. However, rather than splitting the data at a single time point into a training and testing set, we trained the classifier on the data from a given time point and then tested the trained classifier on a different time point^40^. Therefore, the main diagonal (train and test timepoint are the same) corresponds to the classifiers’ training accuracy; the off-diagonal results refer to the classifiers’ test accuracy.

### Simulating Ramping Spike-Trains

We simulated spike trains where we knew that the ground-truth was a gradual ramp in order to see if the asymmetry we observed in our temporal generalization analysis (where decoders trained to differentiate activity from a baseline generalize better into the future than into the past) is due to ramping. We simulated 50 spike trains for a pseudopopulation of 20 neurons. Half of the neurons increased their firing rates from 5 Hz to 8 Hz, and the other half decreased their firing rates from 8 Hz to 5 Hz. We simulated 3 s of activity (labeled as −3 to 0 to match other analyses) and generated spike trains in 50 ms bins, where the instantaneous firing rate was obtained by linearly interpolating the starting and ending firing rates. Then, we conducted the same temporal generalization analysis as above using a baseline of [−3,−2.6]. We found a striking asymmetry in the AUCs such that decoders generalized better into the future than into the past (one-tailed rank-sum test on upper vs lower triangle of AUC matrix, p < 0.001; Fig S4D).

### Analysis of Pairwise Correlations at Baseline

Our model suggests that early ramping reflects fluctuations in clustered networks aligned to threshold-crossings. Crucially, those fluctuations emerge naturally due to the clustered connectivity, and also because neurons that ramp up tend to be in the same cluster, and those that ramp down tend to belong to clusters other than the one ramping up. However, in our model, connectivity is static, as would be expected in biological networks at the time scales in which we simulate here. Neurons in the same cluster therefore show correlated activity across the entire trial, not just during ramping. Hence, our model predicts that neurons that ramp in up or down together before movement or threshold crossings should have correlated activity even before ramping onset.

We thus investigated pairwise correlations at baseline in empirical neurons in a baseline period of (−3.5,−2.5) (a baseline of (−3, −2.5) was used for simulated data because we did not save data earlier than −3 s). We first smoothed (Gaussian kernel, 400 ms wide), then soft-normalized (divided by the square root of the maximum firing rate^39^) firing rates, and finally down-sampled to a sample every 50 ms to avoid inflating correlations due to autocorrelations in the data. After that, we correlated firing rates between every pair of neurons that were recorded in the same session, keeping track of whether the pair was both increasing, both decreasing, one increasing and one decreasing, or involved a neuron that didn’t change its firing rate (for these analyses we categorized neurons with a steplike response as “other” in order to focus solely on slow-ramping behavior). At that point, we used linear mixed-effects models (LME; with the network or subject as a random intercept) to investigate whether correlation structure at baseline depended on later ramping behavior. We found that increasing and decreasing pairs show significantly correlated activity at baseline. In contrast, increasing-decreasing pairs do not; they also show significantly lower correlations than increasing-increasing pairs and decreasing-decreasing pairs. Notably, running the same analysis without separating slow ramping and steplike responses led to all shown statistical tests in Fig. 4C becoming highly significant (both increasing vs increasing-decreasing p = 0.001; both increasing vs other p = 0.013; both decreasing vs increasing-decreasing p < 0.001; both decreasing vs other p < 0.001; LME post-hoc tests, Tukey correction), as well as the difference between increasing-decreasing and other pairs becoming significant (p = 0.040). Additionally, omitting pairwise comparisons involving two neurons on the same channel (accounting for potential issues due to spike sorting) largely did not change the results for decreasing-decreasing pairs, which had many examples. However, only few slow ramping increasing-increasing pairs remained after this correction, reducing statistical power and leading to insignificant differences between increasing-increasing vs increasing-decreasing pairs (p = 0.439), and increasing-increasing vs other pairs (p = 0.777). However, the estimated correlation was similar to decreasing neurons after that correction, so we suggest that the lack of statistical power is the main cause of these differences.

## Supplementary Material

**Supplementary Figure 1:**
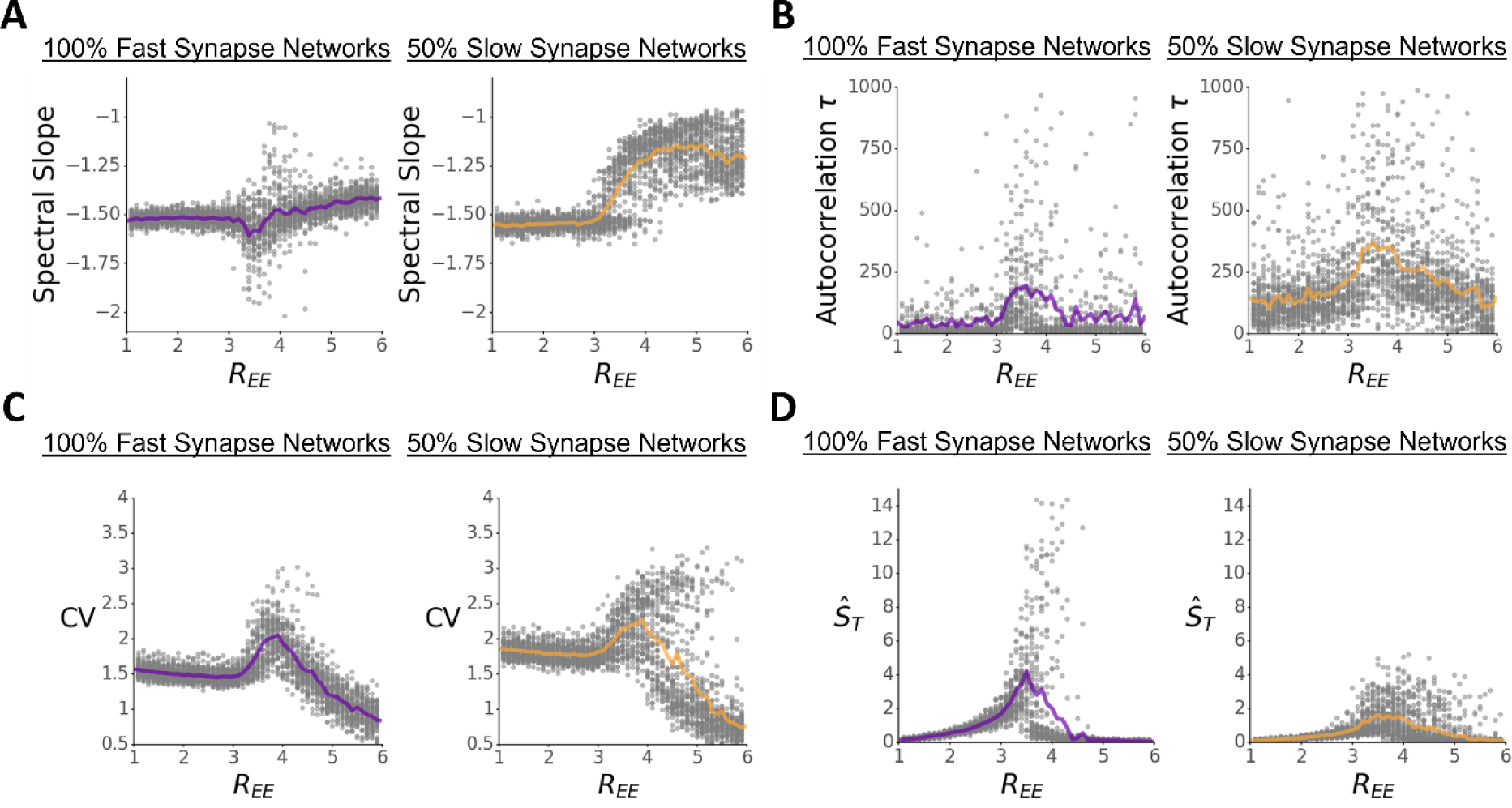
Homogeneity of fluctuation statistics across networks. In all figures, shown are average statistics (solid line), 95% confidence interval (shaded region) and example values for different networks at different REE values. **A:** Spectral slope, as in Fig. 2C. **B:** Autocorrelation timescales, as in Fig. 2D. **C:** Coefficient of variation, as in Fig. 2E. **D:** ŜT measure of slow-switching spiking variability, as in Fig. 2F.

**Supplementary Figure 2:**
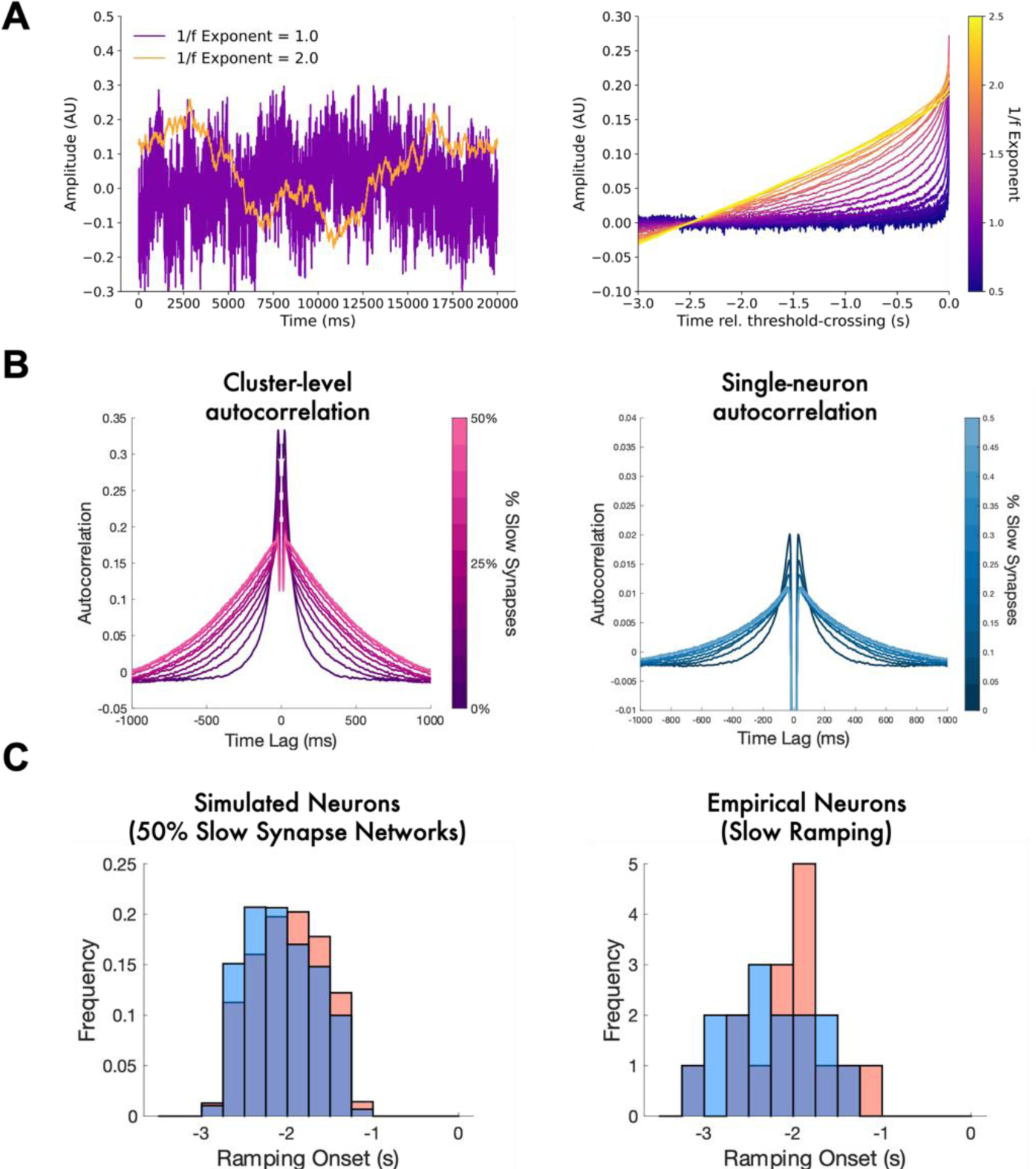
Relation between autocorrelation and ramping onset & characterization of ramping onset in simulated & empirical data. **A:** Relation between autocorrelation and slow ramping in pink noise. We simulated pink noise with 1/f exponents ranging from 0.5 to 2.5 using the *colorednoise* Python package. We simulated 10,000 trials of 20 s of noise and aligned to threshold-crossings (threshold of 0.3, baselined using the period [−3, −2] as in our other simulations). Left: Two examples of simulated trials with lower and higher autocorrelation (1/f exponent of 1 and 2, respectively). Right: When aligned to threshold crossings, high-autocorrelated noise shows slow ramping in the preceding signal, whereas low-autocorrelated noise shows little to no ramping. As autocorrelation (1/f exponent) increases, slow ramping behavior emerges and, at least to some degree, has earlier onsets. **B:** Autocorrelation at the cluster-level (left, autocorrelation calculated for each cluster-average firing rate and then averaged across clusters) and single-neuron level (right, autocorrelation calculated for each neuron’s firing rate and then averaged across neurons). Autocorrelation increases with increased proportion of slow synapses. Note from the Y-Axis that cluster-level autocorrelation is much larger than the single-neuron level. **C:** Characterization of the onset of slow ramping simulated (left) and empirical neurons (right) (red = increasing neurons, blue = decreasing neurons) with respect to movement onset (at t = 0 s). We used the MATLAB function findchangepts with the linear statistical method to conduct a breakpoint analysis in the timeframe of −3.5 to −1 s to determine the onset of ramping. (Determining the onset of a change in slope in a noisy signal is a difficult problem, and methods using means are often biased; hence we used the linear statistical method, which takes into account both mean and slope to find a breakpoint.) In simulated and empirical data, a range of ramping onsets are apparent, with an average ramping onset of about −2 s.

**Supplementary Figure 3:**
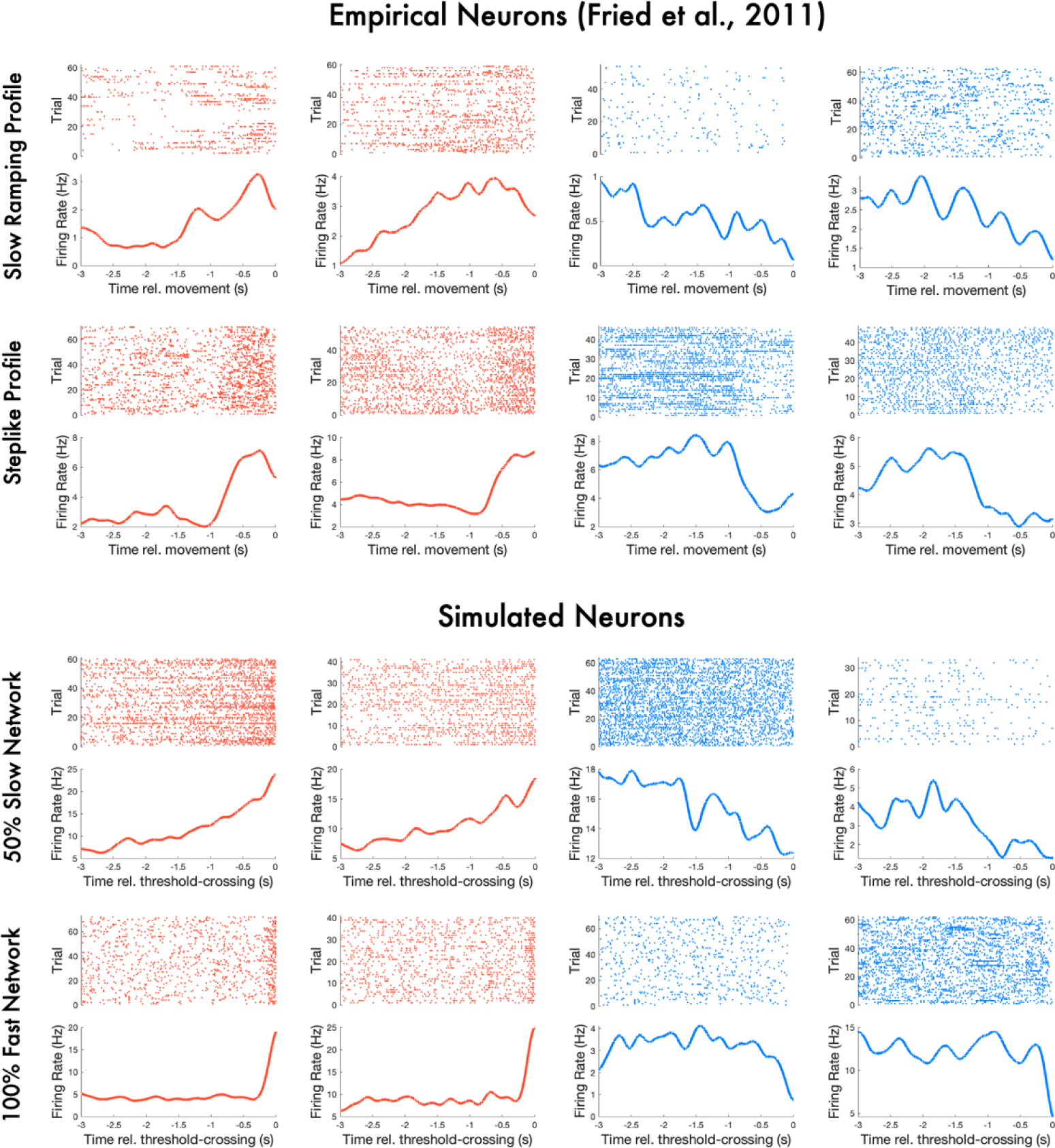
Further example of raster plots & trial-averaged firing rates for empirical and simulated neurons (increasing in red, decreasing in blue).

**Supplementary Figure 4:**
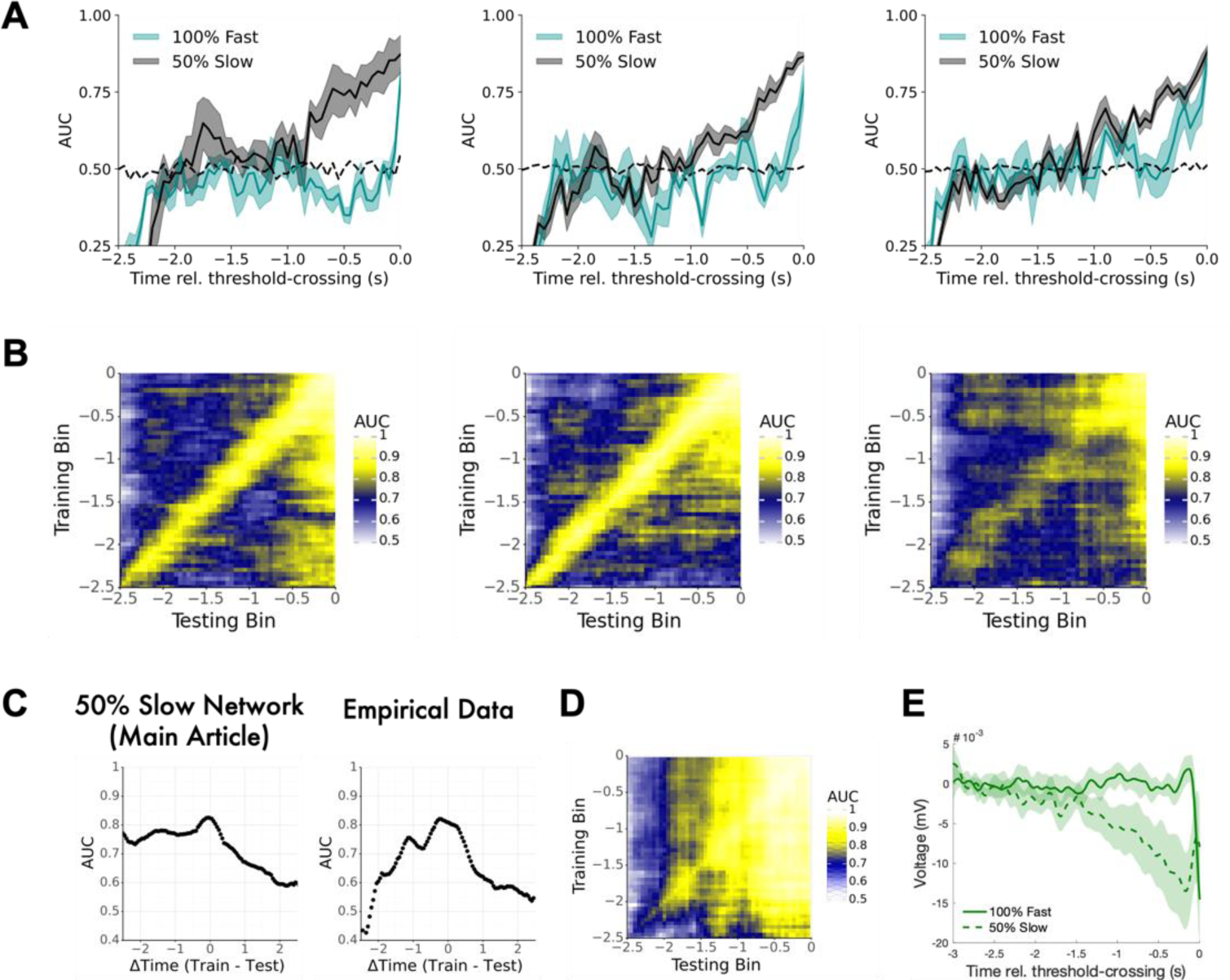
More examples of slow ramping decoding accuracy in simulated networks. **A:** As in Fig. 3A, decoding AUC over time for 3 other example networks with 100% fast (blue) and 50% slow (black) synapses. **B:** As in Fig. 3B, temporal generalization of decoders trained at one time bin and tested on another time bin. **C:** AUC averaged across train-test bin pairs that are the same distance from diagonal. AUC was higher when the training bin was earlier than the test bin (i.e., for negative x-axis values) for both simulated (left) and empirical data (right). That asymmetry is consistent with a largely monotonic drift in population firing rates away from a baseline. **D:** Results of temporal generalization analysis conducted on Poisson spike-trains that ramp. In a case where ramping is a priori known to be present, we see an asymmetry such that decoders that classify activity at a given time from baseline (−3 to −2.6 s in this case) generalize better to the future than they do to the past, indicated by higher AUC in the lower-triangle compared to the upper-triangle of this matrix. **E:** Pre-threshold changes in total PSP delivered to all network neurons (EEG proxy). Lines & shaded region is grand-average mean & 95% confidence intervals across 20 simulated networks. Magnitude is smaller than the EEG proxy used in main text (PSPs onto excitatory neurons only), but still shows an early negative deflection reminiscent of the EEG readiness potential before self-initiated actions.

## Notes

### Competing Interest Statement

The authors have declared no competing interest.

### Summary of Updates

We conducted new analyses resulting in a new main figure (now Figure 2) and more panels in several figures, expanded the methods and discussion.

